# Maternal n-3 PUFA deficiency alters brain fatty acid and oxylipin profiles across perinatal development in offspring

**DOI:** 10.1101/2025.11.02.686109

**Authors:** Marchaland Flore, Martinat Maud, Di Miceli Mathieu, Grégoire Stéphane, Rossitto Moïra, Morel Lydie, Aubert Agnès, Séré Alexandra, Delpech Jean-Christophe, Joffre Corinne, Acar Niyazi, Bazinet Richard P, Layé Sophie

## Abstract

Long-chain polyunsaturated fatty acids (LC-PUFAs), particularly arachidonic acid (AA, 20:4n-6) and docosahexaenoic acid (DHA, 22:6n-3), are essential for optimal neurodevelopment through their effect on neuronal proliferation, neurite outgrowth and synaptogenesis. Emerging evidence highlights that brain PUFAs are metabolized in oxylipins, the bioactive oxidized PUFA metabolites known to regulate inflammatory processes. Recent data highlighted that both PUFA and oxylipin profiles are modulated in the adult male brain by dietary PUFA content. However, little is known on the impact of maternal dietary n-3 PUFA intake during the perinatal period and the neurodevelopmental profile of brain fatty acids and associated oxylipins in offspring, and whether these effects differ between sexes. To address this question, we first measured fatty acid levels in the placenta and embryonic brain of male and female offspring of mothers fed a sufficient or deficient diet in n-3 PUFAs at embryonic day (E)17.5. Then, fatty acids and oxylipins were measured at different post-natal stages, in the brain at P0 and P7, and in the hippocampus at P14 and P21, in both male and female offspring. Our results show that maternal n-3 PUFA dietary deficiency alters fatty acid profiles as early as E17.5 in both the placenta and the brain. Furthermore, dietary intervention affects both fatty acid and oxylipin profiles throughout postnatal brain development, with notable sex-specific differences. These findings underscore the critical importance of adequate maternal n-3 PUFA intake during the perinatal period for maintaining an optimal PUFA and oxylipin profiles, with potential implications for fetal and postnatal brain development.

## Introduction

The brain is highly enriched in fatty acids, including saturated (SFA), monounsaturated (MUFA) and polyunsaturated fatty acids (PUFAs)^1^, which are mainly delivered to the brain *via* the bloodstream following dietary intake^2^. Grey matter is particularly enriched in n-6 and n-3 long-chain PUFAs (LC-PUFAs)^3^, with arachidonic acid (AA, 20:4n-6) and docosahexaenoic acid (DHA, 22:6n-3) being the predominant species esterified on phospholipids of neuronal and glial cell membranes^1, 4, 5^, while eicosapentaenoic acid (EPA, 20:5n-3) is mainly incorporated into microglial membranes^6, 7^. In mammals, AA, EPA and DHA can be synthesized from dietary precursors found in plant sources, namely linoleic acid (LA, 18:2n-6) and α-linolenic acid (ALA, 18:3n-3). These LC-PUFAs can also be directly provided in their preformed state, through the consumption of terrestrial animal products or seafood^8, 9^. Overall, both dietary precursors and preformed LC-PUFAs contribute directly to the levels of LC-PUFAs in the brain^1^. Importantly, the highest accumulation of LC-PUFAs in the brain occurs during the perinatal period. In humans, this accretion is particularly intense from the third trimester of gestation to two years of age^10, 11, 12^, and in rodents, from late gestation to the end of the third postnatal week^13^. During this perinatal period, LC-PUFAs are initially transferred to the embryo *via* the placenta and subsequently to the offspring through breast milk, including delivery to the developing brain^14, 15^. DHA is essential for brain neurodevelopment, as it regulates neuronal proliferation, neurite outgrowth and synaptogenesis, as demonstrated both *in vitro* in rodent neuronal cultures and human iPSC-derived neurons as well as *in vivo*^16, 17, 18, 19, 20, 21, 22^. Overall, a deficiency in LC-PUFAs and/or an imbalance between n-3 and n-6 PUFAs, can impair neurodevelopment^14, 15^. Premature birth or cesarean delivery disrupts materno-fetal LC-PUFA transfer, leading to reduced availability compared to full-term infants, which may compromise neurodevelopment^23, 24^. Moreover, low-birth-weight infants have reduced adipose stores of LC-PUFAs and are consequently at a higher risk of neurodevelopmental disorders^25^. In rodents, maternal dietary n-3 PUFA deficiency during the perinatal period reduces brain DHA levels by embryonic day 17 (E17)^26^, impairs memory and synaptic plasticity from postnatal day 21 (P21)^21, 27, 28^ and increases offspring vulnerability to the detrimental effects of early-life inflammatory challenges^26, 29^. However, the mechanisms linking altered PUFA profiles to impaired brain development remain poorly understood. An additional key aspect of PUFA function in the brain involves their bioactive metabolites. Oxylipins, which form a major subclass of lipid mediators enzymatically derived *via* cyclooxygenase (COX), lipoxygenase (LOX) and cytochrome P450 (CYP450) pathways, play important signaling roles^30, 31, 32^. They are present in the adult brain, with their levels and profiles strongly influenced by inflammatory stimuli and dietary PUFA status. Indeed, hundreds of oxylipins can be synthesized from a limited set of PUFA precursors, mainly DHA, EPA and AA^32, 33, 34, 35, 36, 37^. This is particularly important since oxylipins derived from n-6 and n-3 PUFAs are key regulators of numerous brain processes, including synaptic plasticity, neuronal morphology, neurotransmission, cerebral blood flow and neuroinflammation^2, 7, 31, 32, 38, 39, 40, 41^. Notably, oxylipins derived from n-6 PUFAs (e.g., prostaglandins, leukotrienes, thromboxanes, and lipoxins) and those derived from n-3 PUFAs (e.g., specialized pro-resolving mediators, SPMs) often exert opposing effects on inflammation and neuroinflammation^31, 42^. Recent studies, including our own, have shown that, in a context of maternal dietary n-3 PUFA deficiency, AA and its metabolite 12-HETE (12- Hydroxyeicosatetraenoic acid) altered postnatal maturation of neuronal networks *via* their specific effects on the activity of microglia^21^, the brain resident phagocytic immune cells essential for synaptic pruning^43^. This further supports the notion that oxylipins play a crucial role in brain development. These findings highlight the importance of dietary PUFA composition in shaping the brain’s oxylipin profile and potentially its functional outcomes.

Only a few studies have investigated sex differences in the composition of PUFAs and oxylipins in the brain, most of which have focused on adulthood. In rats and mice maintained under physiological conditions and fed a standard diet, brain fatty acid composition, including AA and DHA, as well as brain oxylipin profile did not differ between sexes^44, 45, 46, 47, 48^. However, sex-dependent variations in fatty acid and oxylipin profiles during brain development, and their potential modulation by dietary n-3 and n-6 PUFAs, remain largely unexplored.

In this study, we aimed to investigate how dietary interventions, using isocaloric diets with different LA/ALA ratios, affect the developmental profiles of brain fatty acids and oxylipins in offspring from E17.5 to PND21, taking into account potential sex differences. To this end, we first measured fatty acid levels in the placenta and embryonic brain at E17.5. We then analyzed fatty acids and quantified oxylipins in the brain at P0 and P7 and in the hippocampus at P14 and P21 in both male and female offspring.

## Materials and Methods

### Animals and diets

Animal housing and experimental procedures were performed in accordance with the European Union directive (2010/63/EU) and validated by the ethics committee (approval #2022031116224183). All experiments were conducted using CD1 mice (Janvier Labs, Le Genest-Saint-Isle, France). Animals were housed under standard conditions with controlled temperature (23 ± 1 °C) and humidity (40-50%), and maintained on a 12-hour light/dark cycle (7:30-19:30). Food and water were available *ad libitum*. Eight-week-old male and female mice were mated overnight. The successful mating was confirmed by the presence of a vaginal plug the following morning that was designated as embryonic day 0.5 (E0.5). Right before mating, mice were randomly assigned to isocaloric experimental diets, differing in their LA/ALA ratio, and maintained throughout gestation and lactation, as previously described^21, 26, 28, 49^. These diets, manufactured by the INRAE unit (Jouy-en-Josas, France), contained 5% of fat in the form of sunflower oil, rich in LA (n-3 PUFA-deficient offspring) or in the form of canola oil, rich in ALA (n-3 PUFA- sufficient offspring) (Tables 1 and 2). Pregnant females were housed individually throughout gestation and lactation. To avoid cage effects, pups used at each developmental stage were selected from distinct litters, with each dam contributing a maximum of two litters to the study. In total, for experiments at E17.5, 3 mothers/diet and 2 pups/litter were used. For postnatal experiments, 2 mothers/diet and 2 to 3 pups/litter were included. Sample sizes (number of pups per experiment) are specified in the figure legends.

**Table 1:** Composition of the experimental diets (g/kg diet). Adapted from Joffre et al., 2016. ^a^ For detailed composition, see Table 2. ^b^ Composition (g/kg): sucrose, 110.7; CaCO3, 240; K2HPO4, 215; CaHPO4, 215; MgSO4,7H2O, 100; NaCl, 60; MgO, 40; FeSO4,7H2O, 8; ZnSO4,7H2O, 7; MnSO4,H2O, 2; CuSO4,5H2O, 1; Na2SiO7,3H2O, 0.5; AlK(SO4)2,12H2O, 0.2; K2CrO4, 0.15; NaF, 0.1; NiSO4,6H2O, 0.1; H2BO3, O.1; CoSO4,7H2O, 0.05; KIO3, 0.04; (NH4)6Mo7O24,4H2O, 0.02; LiCl, 0.015; Na2SeO3, 0.015; NH4VO3, 0.01. ^c^ Composition (g/kg): sucrose, 549.45; retinyl acetate, 1; cholecalciferol, 0.25; DL- α-tocopheryl acetate, 20; phylloquinone, 0.1; thiamin HCl, 1; riboflavin, 1; nicotinic acid, 5; calcium pantothenate, 2.5; pyridoxine HCl, 1; biotin, 1; folic acid, 0.2; cyanobalamin, 2.5; choline HCl, 200; DL-methionin, 200; p-aminobenzoic acid, 5; inositol, 10.

| Ingredient | Amount |
| --- | --- |
| Casein | 180 |
| Cornstarch | 460 |
| Sucrose | 230 |
| Cellulose | 20 |
| Fat<sup>a</sup> | 50 |
| Mineral mix<sup>b</sup> | 50 |
| Vitamin mix<sup>c</sup> | 10 |

**Table 2:** Fatty acid composition of the dietary lipids (% of total fatty acids). Adapted from Joffre et al., 2016. SFAs: saturated fatty acids; MUFAs: monounsaturated fatty acids; PUFAs: polyunsaturated fatty acids; LA: linoleic acid; AA: arachidonic acid; ALA: α-linolenic acid; EPA: eicosapentaenoic acid; DHA: docosahexaenoic acid; ND: not detected (under the limit for the detection by gas chromatography, <0.05%).

| **Diet** | **n-3 PUFA-sufficient** | **n-3 PUFA-deficient** |
| --- | --- | --- |
| 16:0 | 22.6 | 6.2 |
| 18:0 | 3.3 | 4.4 |
| Other SFAs | 1.8 | 1.6 |
| **Total SFAs** | **27.7** | **12.2** |
| 16:1n-7 | 0.2 | 0.1 |
| 18:1n-9 | 57.9 | 26.0 |
| 18:1n-7 | 1.5 | 0.9 |
| Other MUFAs | 0.4 | 0.2 |
| **Total MUFAs** | **60.0** | **27.2** |
| 18:2n-6 (LA) | 10.6 | 60.5 |
| 20:4n-6 (AA) | ND | ND |
| **Total n-6 PUFAs** | **10.7** | **60.5** |
| 18:3n-3 (ALA) | 1.6 | 0.1 |
| 18:4n-3 | ND | ND |
| 20:5n-3 (EPA) | ND | ND |
| 22:5n-3 | ND | ND |
| 22:6n-3 (DHA) | ND | ND |
| **Total n-3 PUFAs** | **1.6** | **0.1** |
| **Total PUFAs** | **12.3** | **60.6** |
| **n-6/n-3 PUFA ratio** | **6.7** | **>500** |

### Tissue collection

Placentas and brains from individual embryos were collected at E17.5 for lipid analysis. As the mouse is a multiparous species, each embryo has its own placenta. Samples were collected following maternal anesthesia with Bupaq solution (Centravet), diluted to 1:30 in water for injectable preparations and euthanasia using Euthasol Vet solution (Centravet), diluted to 1:10 in water for injectable preparations.

For each solution, 10 μl per gram of mouse was injected. A tail sample was also collected from each embryo for determination of sex (see below). At later developmental stages, pups were anesthetized with Bupaq solution and euthanized with Euthasol Vet solution, using the same dilutions and injection volumes described above. Brains were then rapidly extracted and either frozen whole (P0 and P7) or processed for hippocampus dissection (P14 and P21). All tissue collections were performed on ice and samples were immediately stored at -80 °C until further processing. Each brain and hippocampal sample was divided into two equal parts: one for lipid analysis, the other for oxylipin quantification.

### Sex identification at E17.5

To determine embryo sex, genomic DNA was extracted from tail samples and used for Rbm31 genotyping (Tunster, 2017). Briefly, tail tissue was digested in 200 µl of 0.05 M NaOH during 10 min at 100 °C. Then, 50 µl of 1 M Tris-HCl (pH 8.0) were added to the mix. After centrifugation for 5 min, the supernatant containing DNA was collected and stored at 4 °C. Two µl of DNA extract were used for each PCR reaction. Each 20 µl PCR reaction mix also contained 5 µl of deoxynucleotide mix, 10 µl of 2X PCR mix (Promega, WI, USA), 0.1 µl of homemade Taq polymerase, and primers (5’- CACCTTAAGAACAAGCCAATACA-3’ and 5’-GGCTTGTCCTGAAAACATTTGG-3’). PCR amplification in the thermocyler was performed using the following program: 2 min at 94 °C, 30 cycles of 20 s at 94 °C, 20 s at 60 °C, 30 s at 68°C, 5 min at 72 °C, then a cool down to 4 °C. Amplified PCR products were run through a 2% agarose gel and visualized using SafeView (Applied Biosystems, Carlsbad, CA) on UV table with gel imager (EBox CX5, Vilber, Collégien, France).

### Lipid analysis

Lipids were extracted from placentas (E17.5), whole brains (E17.5), half-brains (P0 and P7) and half- hippocampi (P14 and P21) at the Eye & Nutrition Research Group laboratory (CSGA, INRAE, Dijon, France) and further processed for fatty acid measurement. Briefly, total lipids were extracted according to the Folch procedure using a chloroform/methanol mixture^50^. The fatty acids were methylated^51^ and the composition of the resulting fatty acid methyl esters (FAMEs) and dimethyl acetals (DMAs) was determined by gas chromatography coupled to flame ionization detection (GC-FID) using a Trace 1310 gas chromatograph (Thermo Scientific, Courtaboeuf, France), equipped with a split/splitless injector, a flame ionization detector, and a CPSIL-88 capillary column (100 m x 0.25 mm internal diameter; 0.20 µm film thickness; Varian, Les Ulis, France). Hydrogen was used as the carrier gas at an inlet pressure of 210 kPa. The temperature program of the oven was set as follows: 60 °C for 5 min, increased to 165 °C at 15 °C/min (held for 1 min), then to 225 °C at 2 °C/min, with a final hold at 225 °C for 7 min. Injector and detector temperatures were set at 250 °C and 280 °C, respectively. FAMEs were identified by comparison with commercial and synthetic standards. The data were computed using the Chromeleon software (version 7.1.10 ES, Thermo Scientific). Results are expressed as the percentage of total FAMEs and DMAs.

### Oxylipin quantification

Oxylipins derived from LA, DGLA, AA, DHA and EPA were isolated from the remaining half-brain or half-hippocampus samples and analyzed using liquid chromatography–tandem mass spectrometry (LC-MS/MS) at the METATOUL platform (MetaboHUB, INSERM UMR 1048, I2MC, Toulouse, France) (Table 3). Analytical procedures were conducted as previously described^52^. Results were expressed as picograms per milligram of protein (pg/mg) and represented as Z-scores for heatmap visualization.

**Table 3:**
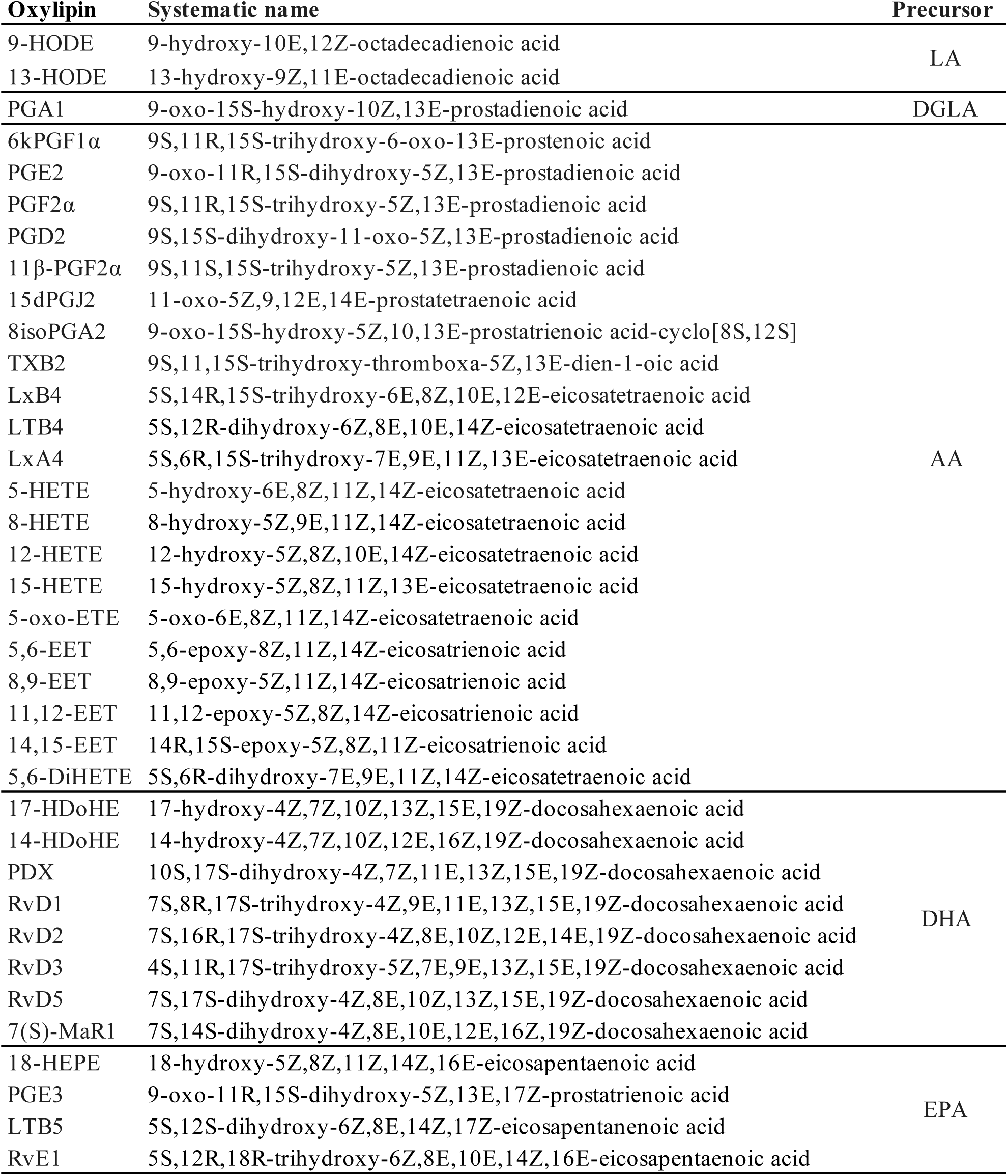
Oxylipins and their PUFA precursors quantified by LC-MS/MS: LA: linoleic acid; DGLA: dihomo-gamma- linoleic acid; AA: arachidonic acid; DHA: docosahexaenoic acid; EPA: eicosapentaenoic acid.

### Statistical analysis

Statistical analyses were performed using GraphPad Prism software (version 10.3.1; GraphPad Software, Boston, MA, USA). Oxylipin and fatty acid levels were analyzed separately at each developmental stage using a two-way ANOVA (sex x diet), followed, when appropriate, by Fisher’s Least Significant Difference (LSD) post hoc tests. Heatmaps of oxylipin levels (expressed as Z-scores) were generated using R software (version 4.4.1) within the RStudio environment (version 2024.09). To allow easier visualization of oxylipin concentrations, data were standardized across samples separately for each sex and developmental stage by computing Z-scores for each individual oxylipin, as previously described (Madore et al., 2020). This normalization involved centering the concentrations around the mean and dividing by the standard deviation, resulting in values with a mean of zero and a standard deviation of one for each oxylipin.

## Results

### 1) Early-life exposure to diets differing in their LA/ALA ratio altered placental and embryo brain PUFA profiles

The impact of maternal n-3 PUFA dietary intervention was first assessed on fatty acid composition of the placenta and embryonic brain at E17.5. Among fatty acids measured, PUFAs were the most affected by early-life exposure to an n-3 PUFA-deficient diet in the placenta (Supplementary Table 1) and brain (Supplementary Table 2) of both males and females. In particular, n-6 PUFAs were significantly increased (Fig. 1A; LA, 20:2n-6, AA, 22:4n-6 and 22:5n-6) while n-3 PUFAs significantly decreased (Fig. 1B; ALA, EPA, 22:5n-3, DHA) in the placenta of n-3 PUFA-deficient males as compared to n-3 PUFA- sufficient males. Accordingly, in the brain, n-6 PUFAs were significantly increased (Fig. 1G; LA, 20:3n- 6, AA, 22:4n-6, 22:5n-6) and n-3 PUFAs decreased (Fig. 1H; EPA, 22:5n-3, DHA) in n-3 PUFA-deficient males as compared to n-3 PUFA-sufficient males. Overall, early-life exposure to an n-3 PUFA-deficient diet similarly affected female placenta and brain, with significantly increased n-6 PUFAs (Fig. 1D for placenta; LA, 18:3n-6, 20:2n-6, 20:3n-6, 22:4n-6, 22:5n-6; Fig. 1J for brain; LA, 20:3n-6, AA, 22:5n-6) and significantly decreased n-3 PUFAs (Fig. 1E for placenta; ALA, EPA, 22:5n-3, DHA; Fig. 1K for brain; EPA, 22:5n-3, DHA) in n-3 PUFA-deficient females as compared to n-3 PUFA-sufficient females. Of note, EPA was significantly higher in the brain of n-3 PUFA-deficient females as compared to n-3 PUFA-deficient males (Supplementary Table 2). Overall, the n-6/n-3 PUFA ratio was significantly increased in the placenta and brain of male (Fig. 1C, 1I) and female (Fig. 1F, 1L) embryos from dams fed an n-3 PUFA-deficient diet as compared to an n-3 PUFA sufficient diet.

**Figure 1:**
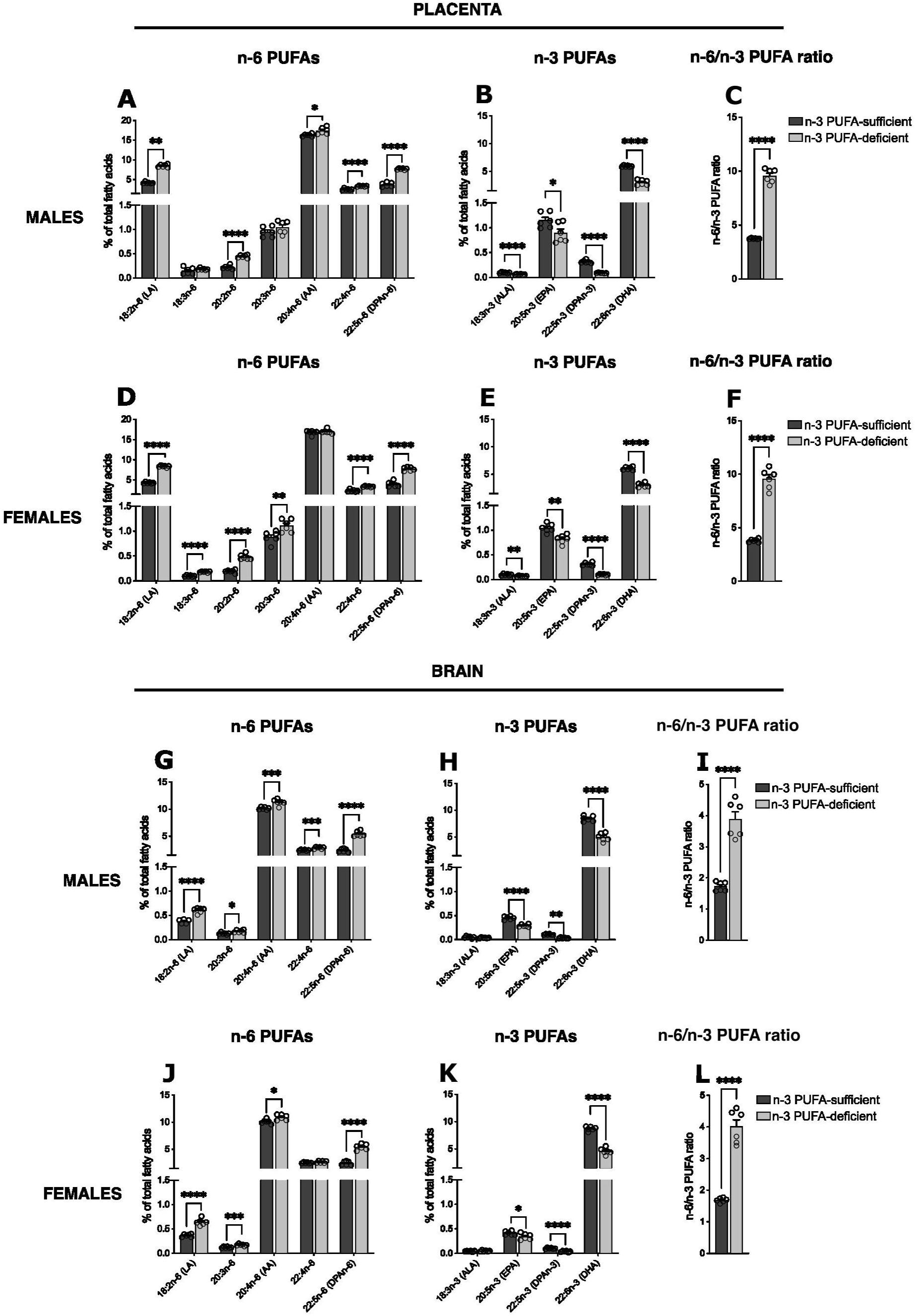
Placental and brain PUFA profiles in male and female embryos: (A-C) Placental levels of n-6 PUFAs (A), n-3 PUFAs (B) and n-6/n-3 PUFA ratio (C) in male embryos. (D-F) Placental levels of n-6 PUFAs (D), n-3 PUFAs (E) and n- 6/n-3 PUFA ratio (F) in female embryos. (G-I) Brain levels of n-6 PUFAs (G), n-3 PUFAs (H) and n-6/n-3 PUFA ratio (I) in male embryos. (J-L) Brain levels of n-6 PUFAs (J), n-3 PUFAs (K) and n-6/n-3 PUFA ratio (L) in female embryos. Data are represented as mean ± SEM (% of total FAMEs and DMAs). N = 6 mice/group. Statistical comparisons were performed for each fatty acid using a 2 way ANOVA (sex x diet), followed by Fisher’s LSD test in case of significant interaction. * p < 0.05; ** p < 0.01; *** p < 0.001; **** p < 0.0001.

### 2) Early-life exposure to diets differing in their LA/ALA ratio altered brain PUFA and oxylipin profiles at birth

Fatty acid and oxylipin profiles were then determined in the brain of offspring at birth. Among fatty acids, PUFAs were the most impacted by early-life exposure to an n-3 PUFA-deficient diet (Supplementary Table 3). Especially, n-6 PUFAs were significantly increased in males (Fig. 2A; LA, 22:5n-6) and females (Fig. 2D; LA, 20:2n-6, 22:5n-6), while n-3 PUFAs were significantly decreased in males (Fig. 2B; EPA, 22:5n-3, DHA) and females (Fig. 2E; EPA, DHA), resulting in a significant increased n-6/n-3 PUFA ratio (Fig. 2C, 2F). Of note, in n-3 PUFA-deficient brains, 20:2n-6 was significantly higher in females, whereas 22:5n-6 and total n-6 PUFAs were significantly lower, as compared to males (Supplementary Table 3).

**Figure 2:**
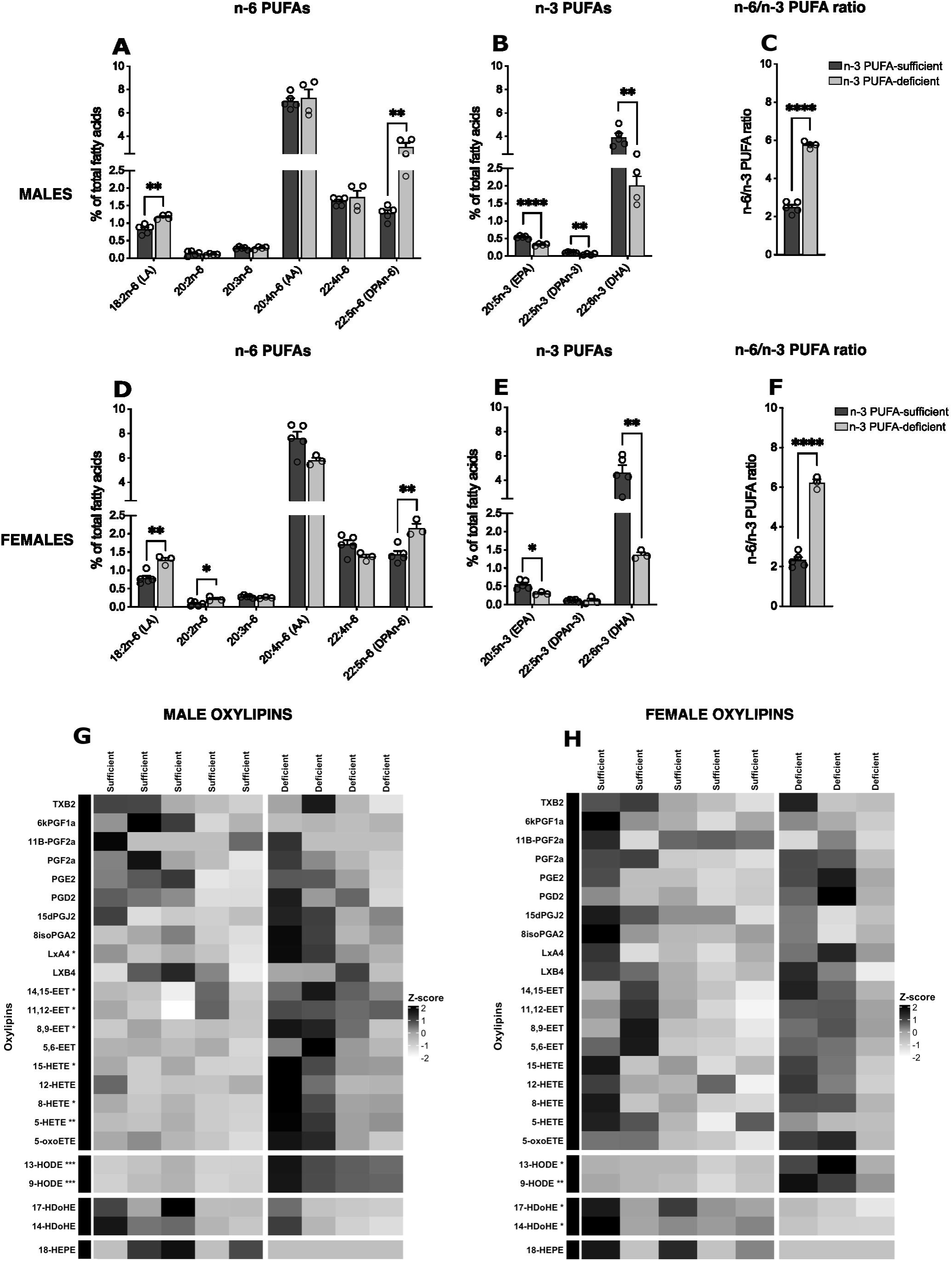
Brain PUFA and oxylipin profiles in the brain of male and female offspring at birth: (A-C) Brain levels of n-6 PUFAs (A), n-3 PUFAs (B) and n-6/n-3 PUFA ratio (C) in male offspring. (D-F) Brain levels of n-6 PUFAs (D), n-3 PUFAs (E) and n-6/n-3 PUFA ratio (F) in female offspring. Data are represented as mean ± SEM (% of total FAMEs and DMAs). N = 3-5 mice/group. (G, H) Heatmaps of oxylipin profiles in the brain of male (G) and female (H) offspring expressed as Z-scores. Rows represent individual oxylipins and columns individual samples. Oxylipins are annotated by their PUFA precursor. Color scale reflects relative abundance (red = higher; blue = lower). Asterisks indicate oxylipins significantly different between groups. Statistical comparisons were performed for each fatty acid and each oxylipin using a 2 way ANOVA (sex x diet), followed by Fisher’s LSD test in case of significant interaction. * p < 0.05; ** p < 0.01; *** p < 0.001; **** p < 0.0001.

Furthermore, oxylipins levels were altered by early-life exposure to n-3 PUFA deficiency (Supplementary Table 4). Particularly, LA-derived 9-HODE and 13-HODE were significantly increased in the brain of males (Fig. 2G) and females (Fig. 2H). Moreover, AA-derived oxylipins LxA4, 14,15-EET, 11,12-EET, 8,9-EET, 15-HETE, 8-HETE and 5-HETE were significantly increased by the n-3 PUFA-deficient diet in the brain of males, but not in females (Fig. 2G). Of note, in n-3 PUFA-sufficient brains, 5-HETE was significantly higher in females than in males (Supplementary Table 4). Additionally, DHA-derived 17- HDoHE and 14-HDoHE were significantly reduced in the brain of n-3 PUFA-deficient females (Fig. 2H).

### 3) Early-life exposure to diets differing in their LA/ALA ratio altered brain PUFA and oxylipin profiles at post-natal week 1

One week after birth, among fatty acids measured, PUFA were the most affected by the n-3 PUFA-deficient diet (Supplementary Table 5). Especially, n-6 PUFAs were significantly increased in males (Fig. 3A; LA, 20:2n-6, AA, 22:4n-6, 22:5n-6) and females (Fig. 3D; LA, AA, 22:4n-6, 22:5n-6), while n-3 PUFAs were significantly decreased in males (Fig. 3B; EPA, 22:5n-3, DHA) and females (Fig. 3E; 22:5n- 3, DHA). Overall, this led to a significant increase of the n-6/n-3 PUFA ratio in the brain of n-3 PUFA-deficient males (Fig. 3C) and females (Fig. 3F). Of note, 20:2n-6 was significantly higher in the brain of n-3 PUFA-deficient females as compared to males (Supplementary Table 5). Moreover, early-life exposure to n-3 PUFA deficiency impacted brain oxylipin profile (Supplementary Table 6). In fact, LA-derived 9-HODE and 13-HODE were significantly increased in the brain of males (Fig. 3G) and females (Fig. 2H). In addition, AA-derived 15-HETE was significantly increased in the brain of males (Fig. 3G), while AA-derived 6kPGF1a and 15-HETE were significantly increased by the n-3 PUFA-deficient diet in the brain of females (Fig. 3H). Of note, in n-3 PUFA-deficient brains, AA-derived LxB4 was significantly higher in females than in males (Supplementary Table 6). Concerning n-3 PUFA-derived oxylipins, early-life exposure to n-3 PUFA deficiency significantly decreased levels of DHA-derived 17-HDoHE and 14- HDoHE, but only in the brain of males (Fig. 3G). In addition, levels of both 17-HDoHE and 14-HDoHE were significantly higher in the brain of males compared to females (Supplementary Table 6).

**Figure 3:**
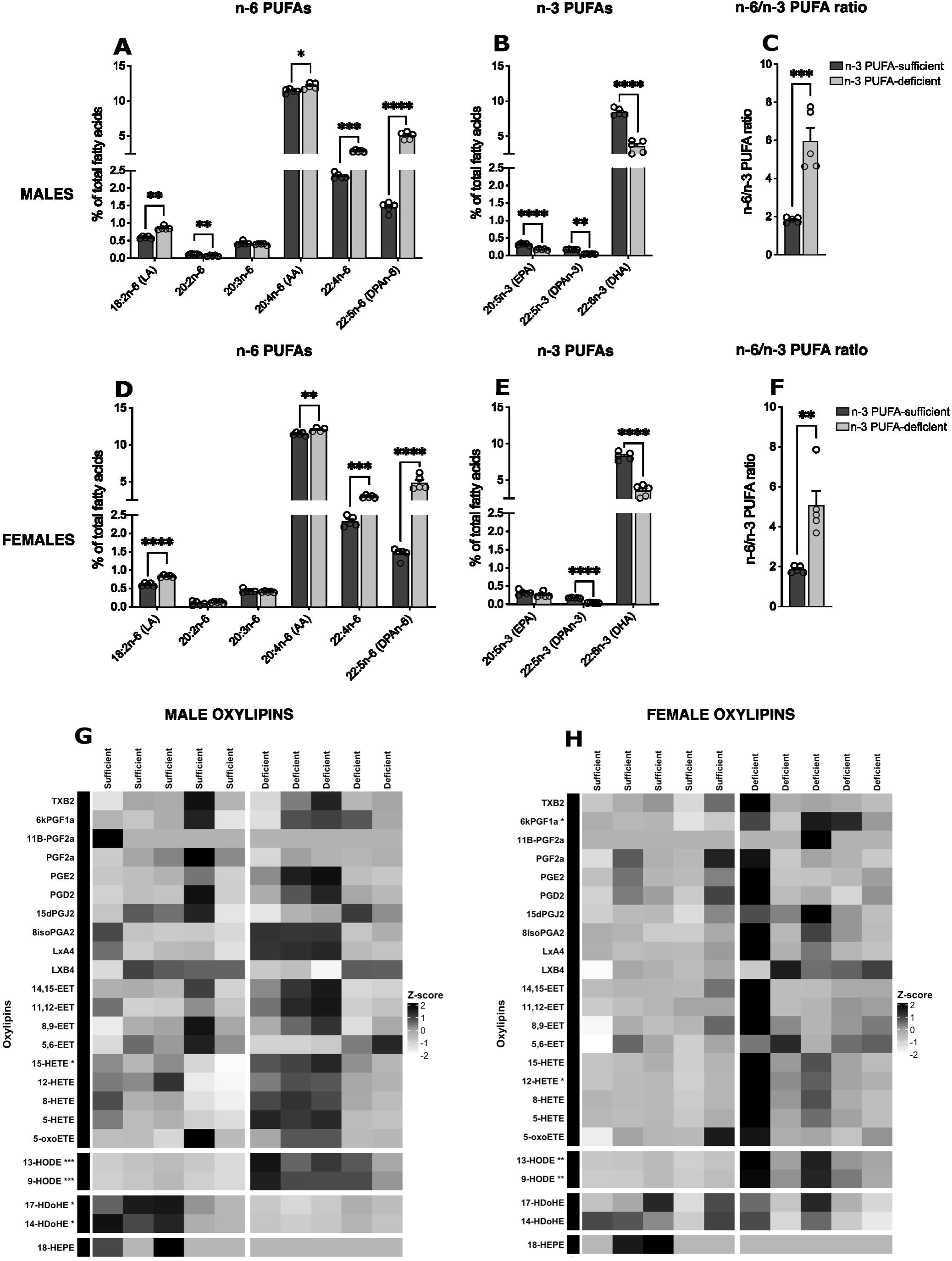
Brain PUFA and oxylipin profiles in the brain of male and female offspring at P7: (A-C) Brain levels of n-6 PUFAs (A), n-3 PUFAs (B) and n-6/n-3 PUFA ratio (C) in male offspring. (D-F) Brain levels of n-6 PUFAs (D), n-3 PUFAs (E) and n-6/n-3 PUFA ratio (F) in female offspring. Data are represented as mean ± SEM (% of total FAMEs and DMAs). N = 5 mice/group. (G, H) Heatmaps of oxylipin profiles in the brain of male (G) and female (H) offspring expressed as Z-scores. Rows represent individual oxylipins and columns individual samples. Oxylipins are annotated by their PUFA precursor. Color scale reflects relative abundance (red = higher; blue = lower). Asterisks indicate oxylipins significantly different between groups. Statistical comparisons were performed for each fatty acid and each oxylipin using a 2 way ANOVA (sex x diet), followed by Fisher’s LSD test in case of significant interaction. * p < 0.05; ** p < 0.01; *** p < 0.001; **** p < 0.0001.

### 4) Early-life exposure to diets differing in their LA/ALA ratio altered hippocampus PUFA and oxylipin profiles at post-natal week 2

Among fatty acids measured in the hippocampus at P14, PUFAs were the most affected by the early-life exposure to an n-3 PUFA-deficient diet (Supplementary Table 7). Accordingly, n-6 PUFAs were significantly increased in males (Fig. 4A; LA, 20:3n-6, AA, 22:4n-6, 22:5n-6) and females (Fig. 4D; LA, 20:3n-6, AA, 22:4n-6, 22:5n-6), while n-3 PUFAs were significantly decreased in males (Fig. 4B; EPA, 22:5n-3, DHA) and females (Fig. 4E; EPA, 22:5n-3, DHA). Consequently, a significant increase of the n- 6/n-3 PUFA ratio was measured in the hippocampus of offspring from n-3 PUFA-deficient mothers (Fig. 4C, 4F). Of note, in n-3 PUFA-sufficient hippocampi, LA was significantly higher in males than in females (Supplementary Table 7). In addition, early-life exposure to n-3 PUFA deficiency altered oxylipin profile in the hippocampus (Supplementary Table 8). Indeed, AA-derived 5,6-EET was significantly increased in n-3 PUFA-deficient females as compared to n-3 PUFA-sufficient females (Fig. 4H).

**Figure 4:**
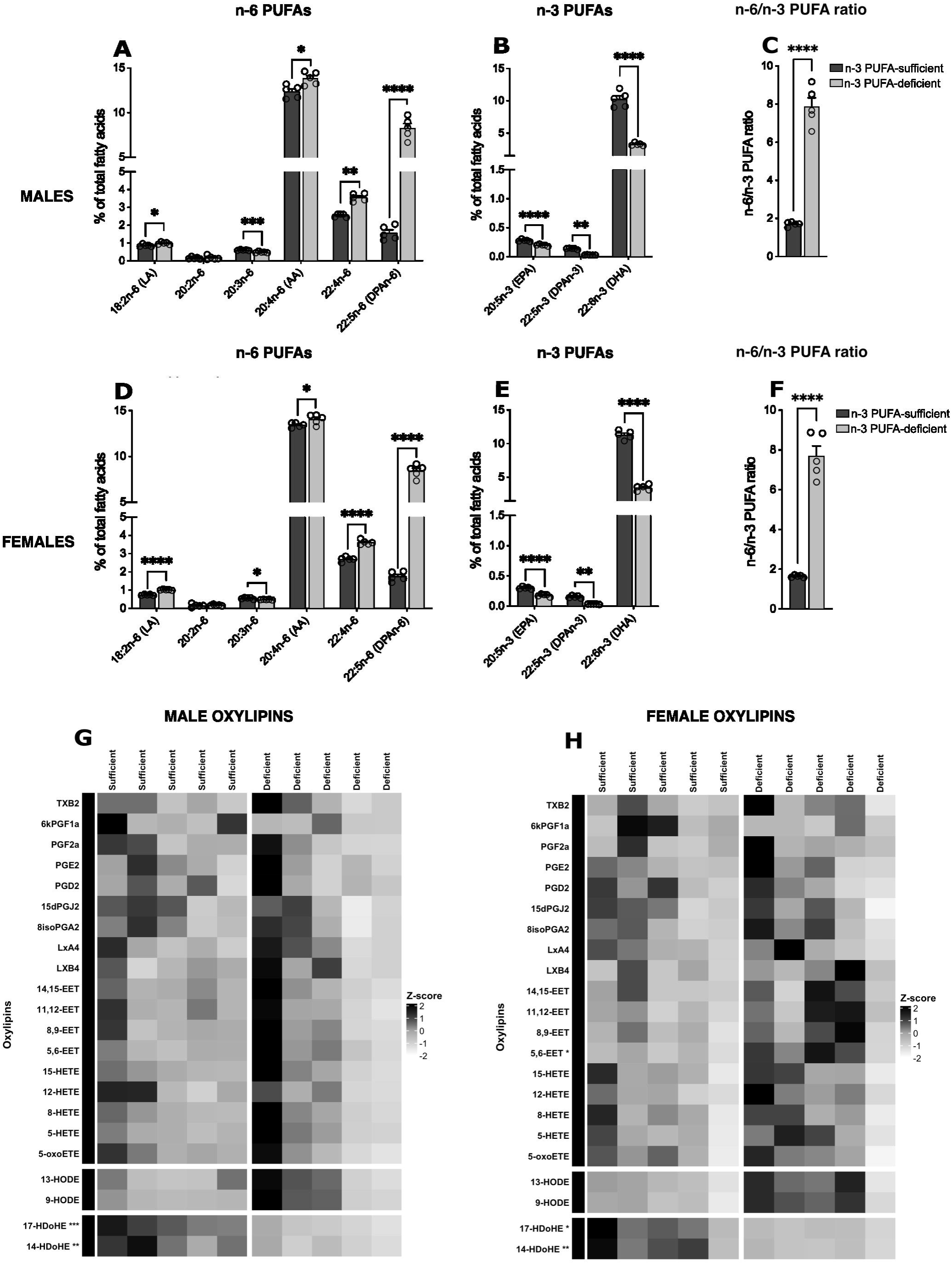
Hippocampal PUFA and oxylipin profiles in the brain of male and female offspring at P14: (A-C) Brain levels of n-6 PUFAs (A), n-3 PUFAs (B) and n-6/n-3 PUFA ratio (C) in male offspring. (D-F) Brain levels of n-6 PUFAs (D), n- 3 PUFAs (E) and n-6/n-3 PUFA ratio (F) in female offspring. Data are represented as mean ± SEM (% of total FAMEs and DMAs). N = 5 mice/group. (G, H) Heatmaps of oxylipin profiles in the hippocampus of male (G) and female (H) offspring expressed as Z-scores. Rows represent individual oxylipins and columns individual samples. Oxylipins are annotated by their PUFA precursor. Color scale reflects relative abundance (red = higher; blue = lower). Asterisks indicate oxylipins significantly different between groups. Statistical comparisons were performed for each fatty acid and each oxylipin using a 2 way ANOVA (sex x diet), followed by Fisher’s LSD test in case of significant interaction. * p < 0.05; ** p < 0.01; *** p < 0.001; **** p < 0.0001.

Moreover, DHA-derived 17-HDoHE and 14-HDoHE were significantly decreased in males (Fig. 4G) and females (Fig. 4H).

### 5) Early-life exposure to diets differing in their LA/ALA ratio altered hippocampus PUFA and oxylipin profiles at post-natal week 3

Finally, among fatty acids measured in the hippocampus at P21, the n-3 PUFA-deficient diet had the most influence on PUFAs (Supplementary Table 9). Indeed, significant increases of n-6 PUFAs were measured in the hippocampi of males (Fig. 5A; LA, 20:3n-6, 22:4n-6, DPAn-6) and females (Fig. 5D; LA, 20:3n-6, 22:4n-6, DPAn-6) as well as significant decreases of n-3 PUFAs in the hippocampus of males (Fig. 5B; EPA, 22:5n-3, DHA) and females (Fig. 5E; EPA, 22:5n-3, DHA). Overall, this led to a significant increase of the n-6/n-3 PUFA ratio in the hippocampus of n-3 PUFA-deficient males (Fig. 5C) and females (Fig. 5F). Additionally, oxylipin profile in the hippocampus was impacted by the n-3 PUFA-deficient diet (Supplementary Table 10). In particular, levels of AA-derived LxA4 were significantly increased in males (Fig. 5G), while those of AA-derived 6kPGF1a were significantly increased in females (Fig. 5J).

**Figure 5:**
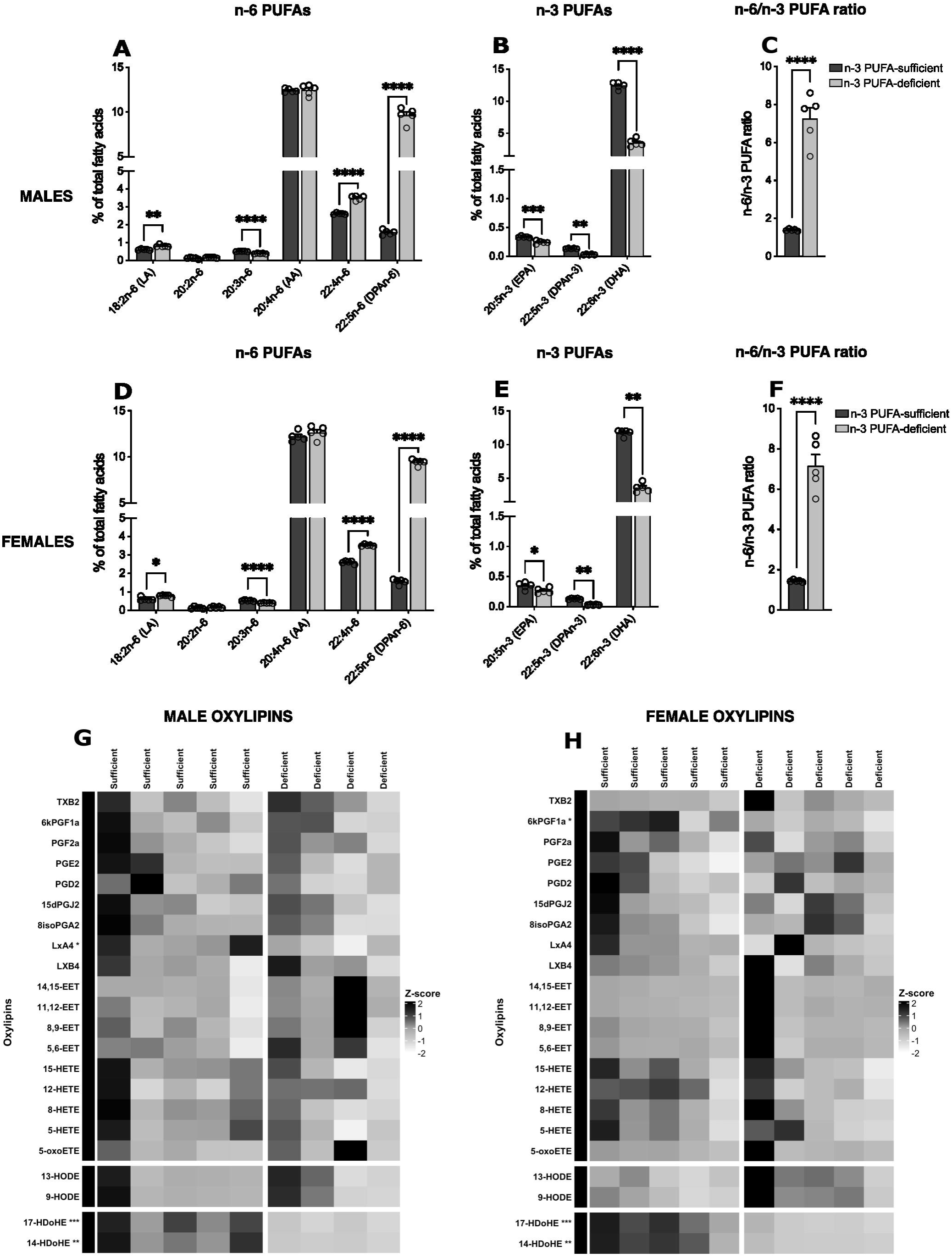
Hippocampal PUFA and oxylipin profiles in the brain of male and female offspring at P21: (A-C) Brain levels of n-6 PUFAs (A), n-3 PUFAs (B) and n-6/n-3 PUFA ratio (C) in male offspring. (D-F) Brain levels of n-6 PUFAs (D), n- 3 PUFAs (E) and n-6/n-3 PUFA ratio (F) in female offspring. Data are represented as mean ± SEM (% of total FAMEs and DMAs). N = 5 mice/group. (G, H) Heatmaps of oxylipin profiles in the hippocampus of male (G) and female (H) offspring expressed as Z-scores. Rows represent individual oxylipins and columns individual samples. Oxylipins are annotated by their PUFA precursor. Color scale reflects relative abundance (red = higher; blue = lower). Asterisks indicate oxylipins significantly different between groups. Statistical comparisons were performed for each fatty acid and each oxylipin using a 2 way ANOVA (sex x diet), followed by Fisher’s LSD test in case of

Moreover, DHA-derived 17-HDoHE and 14-HDoHE were significantly decreased in males (Fig. 5G) and females (Fig. 5H).

## Discussion

In this work, we investigated the impact of perinatal exposure to diets sufficient or deficient in n-3 PUFAs on the fatty acid and their metabolite (oxylipin) profiles in the developing brain of male and female offspring, from embryonic life to weaning, a period during which pups are dependent on maternal PUFA supply. To our knowledge, these are the first data examining the impact of maternal n-3 PUFA dietary intervention on brain PUFA and oxylipin profiles from late embryonic life to postnatal weaning, taking into account sex. Overall, our study brings new insights on the effect of maternal dietary n-3 PUFA intake on postnatal trajectory of offspring brain PUFA, altered as early as E17.5, and oxylipin profiles, impacted from birth, in both males and females.

Maternal PUFAs are crucial for fetal brain development and for optimal feto-placental growth. The placenta plays a central role in shaping the *in utero* environment through gas exchange, nutrient and hormone transport as well as secretion of signaling molecules that regulate both maternal physiology and fetal development. The placenta functions not merely as a passive conduit but actively adapts its transport and signaling in response to maternal and fetal cues, thereby affecting fetal growth trajectories and long-term health outcomes. Disruptions in placental function, due to maternal diet, disease, or suboptimal environmental conditions, can induce metabolic reprogramming and epigenetic changes in the fetus, thereby affecting disease susceptibility later in life^53^. In this study, we reported that at E17.5, an n-3 PUFA-deficient diet initiated at E0 strongly impacted both placenta and brain PUFA profiles, as compared to an n-3 PUFA-sufficient diet. Several n-6 PUFA species were significantly increased, while several n-3 PUFAs were significantly decreased in both placenta and brain following exposure to the n-3 PUFA-deficient diet, resulting in an overall increase of the n-6 to n-3 PUFA ratio in the placenta and brain of both male and female offspring. Our findings also showed that n-6 PUFAs were predominantly present in the placenta, with LA and AA displaying the highest levels in both male and female placentas (around 4% and 17% respectively in n-3 PUFA-sufficient pups; around 8% and 17% respectively in n-3 PUFA-deficient pups). These concentrations were observed regardless of diet. In contrast, n-3 PUFA levels were lower, with DHA being the predominant n-3 PUFA (about 6% in n-3 PUFA-sufficient pups; about 3% in n-3 PUFA-deficient pups), while ALA was detected at the lowest levels (around 0.1% in n-3 PUFA- sufficient pups; around 0.07% in n-3 PUFA-deficient pups). These results are in accordance with our previous study reporting modulation of fatty acid content in the placenta and fetal brain according to the diet content in n-3 PUFAs^26^. They are also in accordance with other reports describing such differences in placental fatty acid profiles in mouse^54, 55^ and human^56^ according to the diet. However, these previous analyses did not distinguish between males and females. Here, we showed that the same PUFA species were affected in both the placenta and brain of male and female pups. Only EPA levels were differentially impacted by sex in the brain of n-3 PUFA-deficient pups on E17.5 (Supplementary Table 2). The majority of LC-PUFAs transported from maternal blood across the placenta to the embryo, including its brain, originate from the maternal liver, although adipose tissue can also contribute^57^. Maternal PUFAs are transferred to the fetus exclusively *via* the placenta, facilitated by specialized fatty acid transporters including FATPs, FAT/CD36, FABPpm as well as Mfsd2a which contributes to the unidirectional transfer of DHA from the mother to the fetus^58, 59^. More recently, a specific transcriptional program in the maternal liver, that promotes the esterification of DHA into phospholipids, has been identified as an additional mechanism supplying LC-PUFAs to the embryo, complementing the dietary LC-PUFAs derived from triglycerides and cholesterol esters^60^. However, despite the activity of this endogenous synthetic pathway in the maternal liver during pregnancy, a decrease in maternal dietary n-3 PUFAs strongly impacts fetal brain DHA content, as seen in the present study and in previous studies^26, 61^. This early decrease in brain DHA, accompanied by a compensatory increase in DPAn-6, starting as soon as 17 days of exposure to the n-3 PUFA-deficient diet, persists throughout postnatal life^21, 49^, including adulthood^1, 26^ , as we previously described in male mice. Here, we further showed that it also occurs in the female brain (see Fig. 2 to 5).

Overall, the same PUFA species were affected in the brains of both males and females, with an increase in n-6 PUFAs and a decrease in n-3 PUFAs under the n-3 PUFA-deficient diet. Only a few PUFA levels were significantly influenced by sex during postnatal development under n-3 PUFA deficiency: 20:2n-6 levels were approximately two-fold higher in females than in males at birth and P7, whereas DPAn-6 levels were around 1.4-fold higher in males than in females at birth (see Supplementary Tables 3 and 5).

To our knowledge, no sex differences in brain PUFA levels have been reported in rodents^46, 47^, suggesting that the limited sex effects observed here across postnatal development may be revealed by dietary n-3 PUFA deficiency. Altogether, our results are consistent with previous findings showing that increased maternal dietary intake of n-3 PUFAs upregulates placental genes involved in fatty acid transport, such as Mfsd2a, thereby enhancing DHA transfer to the fetal brain and promoting neurotrophic signaling. In addition, a maternal diet high in n-3 PUFAs increases both placental and fetal brain Mfsd2a mRNA expression, correlating with higher DHA accumulation in fetal brain^62^. Moreover, alterations in the expression of placental fatty acid metabolism and transport proteins have been observed in mice under conditions of maternal n-3 PUFA deficiency^54, 55^, as well as in humans in relation to maternal metabolic status, such as obesity^63^. Therefore, a few studies measuring PUFA transporters in the placenta and embryonic brain would be of interest.

We also measured oxylipin profiles derived from n-6 (LA, AA) and n-3 (DHA, EPA) PUFAs in the offspring’s brain at P0 and P7 and in the hippocampus at P14 and P21 of males and females from mothers fed a diet low or high in n-3 PUFAs, using a targeted oxylipidome approach, with results presented as Z-scores. Interestingly, oxylipin profiles displayed some age- and sex-related differences, not attributable to dissection duration known to influence brain oxylipin concentrations^64^. Our results particularly pinpoint that 9-HODE and 13-HODE, LA-derived oxylipins previously identified in the adult brain^35, 47^, were upregulated in the brain at P0 and P7 in both males and females, but not in the hippocampus at P14 and P21. These oxidized LA metabolites are produced enzymatically by LOX enzymes and belong to the OXLAMs, a class of oxidized LA derivatives generated through the enzymatic activity of LOX, CYP450, and soluble epoxide hydrolase (sEH)^65, 66, 67, 68^. OXLAMs have also been detected in the developing brain, where they represent the most abundant oxylipins compared to those derived from other PUFAs. This specific pattern is no longer observed in the adult brain^39^. Among OXLAMs, 13-HODE is the most abundant, followed by 9-HODE^38, 39, 69^. Interestingly, both 9-HODE and 13-HODE have been shown to modulate the morphogenesis of rat cortical neurons *in vitro* in a sex-dependent manner, preferentially affecting dendritic and axonal growth while not affecting synapse formation^39, 41^. 13-HODE has also been shown to modulate neurotransmission and synaptic function in the brain^38^. In addition, 9-HODE and 13- HODE bind to lipid receptors such as PPARγ and G2A (GPR132), which are expressed on nociceptive neurons^70, 71^. GPR132 has also been implicated in the 9-HODE-induced increase in astrocyte migration^72^. Notably, recent studies indicate that low plasma levels of 9-HODE predict the severity of brain damage in patients with multiple sclerosis^73^. Altogether, these findings suggest that the LA-derived oxylipins 9- HODE and 13-HODE contribute to neuronal network establishment and exert neuroprotective effects, in addition to their known anti-inflammatory and metabolic effects^74^. However, whether their increase in the developing brain, linked to dietary increases in LA, is beneficial for neurodevelopment remains unclear, as previous studies have reported that LA-rich diets impair developmental brain wiring^21^. This question is particularly relevant given that the LA content of infant formulas ranges from 9% to 20% of total fatty acids^75^. Our results also revealed that dietary n-3 PUFA deficiency modulated lipoxin A4 (LxA4) levels in a sex-dependent manner. In males, LxA4 was increased in the brain at birth but decreased in the hippocampus at P21, whereas no significant changes were measured in females. LxA4 is an AA-derived oxylipin involved in the resolution of inflammation, including in the brain^76^, and has also been suggested to act as a signaling molecule regulating axonal and dendritic outgrowth *via* the formyl-peptide receptor 2 (FPR2)^77^. On the other hand, several AA-derived metabolites were increased in the developing male brain, but not in females, particularly at birth. The bioactive species upregulated by maternal dietary n-3 PUFA deficiency included epoxyeicosatrienoic acids (EETs), generated by CYP450 epoxygenases, and hydroxyeicosatetraenoic acids (HETEs), produced by CYP450 hydroxylases and LOX enzymes. HETEs are well known for their inflammatory properties and have been associated with brain disease severity and neurodegenerative processes^78^, while EETs are recognized for their anti-inflammatory and neuroprotective effects^79, 80^. However, the presence and role of these metabolites in the developing brain has been poorly addressed. We previously reported that 12-HETE, which is increased in the developing male brain at weaning under a maternal n-3 PUFA-deficiency, impairs brain wiring and memory through its effects on microglia^21^. Furthermore, it was shown that EETs are produced endogenously by embryonic cortical neurons and stimulate axonal growth^81^. Furthermore, it was shown that EETs, are produced endogenously by embryonic cortical neurons and that they stimulate axonal growth^81^. Additional studies are necessary to further understand the cellular sources and functions of HETEs and EETs, which are increased in the developing male brain by maternal n-3 PUFA-deficient dietary intervention. We also found that DHA-derived metabolites 17-HDoHE and 14-HDoHE were reduced in the n-3 PUFA-deficient brain at P0 and P7 in a sex-dependent manner, being reduced exclusively in females at P0 and exclusively in males at P7. In the hippocampus at P14 and P21, these hydroxydocosahexaenoic acids were decreased in both male and female n-3 PUFA-deficient mice. 14- and 17-HDoHE are derived from DHA *via* the 12- and 15-LOX pathways and have been reported to increase in the brain of adult mice and rats fed a diet rich in n-3 LC-PUFAs, particularly EPA and DHA^35, 82, 83^. This positive association between these hydroxydocosahexaenoic acids and dietary n-3 PUFA levels is consistent with our findings in the developing brain. 17-HDoHE and 14-HDoHE serve as precursors of several SPMs, notably D-series resolvins and maresins, respectively, which has been described as pro-resolving in inflammation^1, 30^, but are rarely detected in the brain^33^. However, whether fluctuations in HDoHE levels reflect neuroinflammatory conditions remains to be determined. These metabolites, along with other AA-derived oxylipins, are also detected in human cerebrospinal fluid and fluctuate in response to neuroinflammatory events such as surgery^84^. However, to our knowledge, the relationship between oxylipin profiles and dietary PUFA intake in humans remains poorly investigated. Interestingly, during pregnancy, DHA-derived oxylipin levels are increased in the blood of women receiving n-3 LC-PUFA supplementation^85^, as well as in the breast milk of lactating women consuming a DHA-rich diet^86^. However, the impact on the offspring’s oxylipin profile remains to be determined.

In conclusion, our data indicate that brain PUFA levels are largely similarly affected in males and females by maternal n-3 PUFA deficiency during embryonic and postnatal development. However, their metabolism into oxylipins is differentially influenced by sex under n-3 PUFA-deficient conditions, particularly early in postnatal development. Our study further underscores the importance of adequate maternal n-3 PUFA intake for maintaining optimal PUFA and oxylipin profiles, and potentially for supporting fetal and postnatal brain development.

## Data availability

All data are contained within the manuscript.

## Acknowledgments

We thank CIRCE, the animal housing facilities of INRAE UMR1286 NutriNeuro, funded by INRAE, GPR BRAIN_2030, and the Université de Bordeaux. We are grateful to all the zootechnicians of UMR1286 NutriNeuro for breeding our mouse colonies. We thank the MetaToul-MetaboHUB Lipidomic platform for oxylipin quantification and analysis, and the CSGA laboratory for fatty acid measurement and analysis.

## Author contributions

FM, MR, MM, LM, AA, AS, SG generated the data; FM, SG, NA, SL analyzed the data together with the other authors; MDM, NA, CJ and RPB critically contributed to the discussion; SL conceptualized all studies and supervised the work; SL and FM wrote the manuscript and all authors edited and approved the final version of the manuscript.

## Declaration of interests

The authors declare no competing interests.

## Notes

**Funding sources:** This work was supported by INRAE Food4BrainHealth; Fondation pour la Recherche sur le Cerveau CONNECT (SL), Fondation pour la Recherche Médicale PRINSS DEQ 20170336724 (SL); Aquitaine Region AUTISME 2019-1R3M08 (SL) and ExoMarquAge 13059720-13062120 (SL, JCD); ANR IBRAA (SL); ANR-21-JPW2-0004-05 SOLID (SL); PhD extension grant FRM FDT202204014903 (MM); Agence Nationale de la Recherche [ANR-11-LABX-0021-01], INRAE, French “Investissements d’Avenir” Program, Conseil Régional Bourgogne, Franche-Comté (PARI grant), the FEDER (European Funding for Regional Economical Development).

### Competing Interest Statement

The authors have declared no competing interest.

