## Supplementary Data for "Maternal n-3 PUFA deficiency alters brain fatty acid and oxylipin profiles across perinatal development in offspring"

| Sex | Male | | Female | | 2 way ANOVA | | |
|---|---|---|---|---|---|---|---|
| Diet | n-3 PUFA-sufficient | n-3 PUFA-deficient | n-3 PUFA-sufficient | n-3 PUFA-deficient | Diet effect | Sex effect | Interaction |
| Fatty acid | Mean ± SEM | Mean ± SEM | Mean ± SEM | Mean ± SEM | | | |
| 14:0 | 0.26±0.01 | 0.26±0.01 | 0.23±0.01 | 0.25±0.01 | 0.4196 | 0.0567 | 0.2174 |
| 15:0 | 0.14±0.01 | 0.12±0.01 | 0.13±0.02 | 0.12±0.01 | 0.2582 | 0.7022 | 0.7022 |
| 16:0 | 18.02±0.19 | 17.71±0.16 | 17.92±0.09 | 17.85±0.27 | 0.3282 | 0.9132 | 0.5349 |
| 17:0 | 0.16±0.01 | 0.20±0.01 | 0.17±0.01 | 0.20±0.01 | *0.0238 | 0.6877 | 0.6877 |
| 18:0 | 21.02±0.14 | 21.92±0.20 | 21.13±0.13 | 22.18±0.29 | ****<0.0001 | 0.3736 | 0.7196 |
| 20:0 | 0.21±0.01 | 0.20±0.01 | 0.20±0.01 | 0.20±0.01 | 0.6229 | 0.3793 | 0.4924 |
| 22:0 | 0.30±0.03 | 0.37±0.03 | 0.27±0.01 | 0.36±0.02 | **0.0032 | 0.4077 | 0.7137 |
| **Total SFAs** | **40.12±0.18** | **40.78±0.16** | **40.05±0.12** | **41.16±0.14** | ******<0.0001** | **0.3126** | **0.1498** |
| DMA 16:0 | 2.17±0.03 | 2.10±0.01 | 2.10±0.04 | 2.14±0.04 | 0.5979 | 0.5637 | 0.0935 |
| DMA 18:0 | 1.13±0.02 | 1.26±0.01 | 1.15±0.03 | 1.33±0.06 | ***0.0001 | 0.2011 | 0.4255 |
| DMA 18:1n-9 | 0.46±0.01 | 0.29±0.01 | 0.46±0.01 | 0.31±0.01 | ****<0.0001 | 0.3534 | 0.3534 |
| DMA 18:1n-7 | 0.21±0.01 | 0.17±0.01 | 0.19±0.01 | 0.18±0.01 | *0.0183 | 0.7012 | 0.2006 |
| **Total DMAs** | **3.97±0.05** | **3.82±0.03** | **3.89±0.05** | **3.95±0.11** | **0.5203** | **0.6847** | **0.1365** |
| 16:1n-9 | 0.44±0.01 | 0.28±0.01 | 0.43±0.01 | 0.29±0.01 | ****<0.0001 | 0.7066 | 0.3326 |
| 16:1n-7 | 0.70±0.03 | 0.65±0.04 | 0.69±0.04 | 0.63±0.06 | 0.2152 | 0.6925 | 0.8357 |
| 18:1n-9 | 13.71±0.17 | 8.69±0.12 | 13.64±0.25 | 8.54±0.27 | ****<0.0001 | 0.603 | 0.8619 |
| 18:1n-7 | 3.05±0.11 | 2.10±0.03 | 2.86±0.09 | 2.12±0.07 | ****<0.0001 | 0.3056 | 0.1943 |
| 20:1n-9 | 0.31±0.02 | 0.19±0.01 | 0.28±0.02 | 0.21±0.01 | ****<0.0001 | 0.6233 | 0.2169 |

| | | | | | | | |
|---|---|---|---|---|---|---|---|
| 22:1n-9 | 0.16±0.00 | 0.09±0.00 | 0.16±0.00 | 0.10±0.00 | ****<0.0001 | 0.1817 | 0.3672 |
| 24:1n-9 | 0.75±0.06 | 0.52±0.05 | 0.60±0.02 | 0.50±0.02 | ***0.0007 | 0.0527 | 0.1309 |
| **Total MUFAs** | **19.11±0.26** | **12.52±0.23** | **18.64±0.36** | **12.38±0.37** | ******<0.0001** | **0.3366** | **0.6139** |
| 18:2n-6 | 4.19±0.09 | 8.50±0.16 | 4.30±0.07 | 8.36±0.09 | ****<0.0001 | 0.8835 | 0.2581 |
| 18:3n-6 | 0.15±0.03 | 0.18±0.01 | 0.10±0.01 | 0.18±0.01 | **0.0011 | 0.0894 | 0.1097 |
| 20:2n-6 | 0.22±0.02 | 0.44±0.01 | 0.19±0.01 | 0.48±0.02 | ****<0.0001 | 0.804 | 0.0775 |
| 20:3n-6 | 0.95±0.04 | 1.04±0.05 | 0.88±0.05 | 1.11±0.05 | **0.0039 | 0.9867 | 0.1855 |
| 20:4n-6 | 16.27±0.13 | 17.42±0.35 | 16.75±0.21 | 16.98±0.21 | **0.0092 | 0.9259 | 0.0708 |
| 22:4n-6 | 2.51±0.09 | 3.32±0.05 | 2.37±0.11 | 3.36±0.06 | ****<0.0001 | 0.5369 | 0.2835 |
| 22:5n-6 | 3.72±0.23 | 7.69±0.09 | 3.96±0.28 | 7.73±0.18 | ****<0.0001 | 0.4976 | 0.6367 |
| **n-6 PUFAs** | **28.00±0.11**[b] | **38.60±0.14**[a] | **28.54±0.27**[b] | **38.20±0.27**[a] | ******<0.0001** | **0.7313** | ***0.0408** |
| 18:3n-3 | 0.10±0.00 | 0.06±0.00 | 0.10±0.01 | 0.07±0.00 | ****<0.0001 | 0.4678 | 0.4678 |
| 20:5n-3 | 1.15±0.06 | 0.89±0.08 | 1.04±0.04 | 0.82±0.04 | ***0.0004 | 0.1313 | 0.7607 |
| 22:5n-3 | 0.32±0.01 | 0.09±0.00 | 0.31±0.01 | 0.10±0.00 | ****<0.0001 | 0.8577 | 0.4758 |
| 22:6n-3 | 5.93±0.06 | 3.01±0.19 | 6.08±0.12 | 3.04±0.16 | ****<0.0001 | 0.5601 | 0.6827 |
| **n-3 PUFAs** | **7.50±0.04** | **4.06±0.12** | **7.53±0.09** | **4.02±0.15** | ******<0.0001** | **0.8918** | **0.8918** |
| **n-6/n-3 PUFA** | **3.74±0.02** | **9.56±0.26** | **3.80±0.06** | **9.56±0.38** | ******<0.0001** | **0.8937** | **0.9104** |
| 20:3n-9 | 1.31±0.10 | 0.22±0.02 | 1.37±0.11 | 0.30±0.08 | ****<0.0001 | 0.4826 | 0.9088 |
| **Total PUFAs** | **36.81±0.21** | **42.87±0.21** | **37.43±0.33** | **42.52±0.24** | ******<0.0001** | **0.5999** | **0.069** |

**<u>Supplementary Table 1:</u>** Placental fatty acid composition at E17.5: Data are represented as mean ± SEM (% of total FAMEs and DMAs). N = 6 mice/group.

Statistical comparisons were performed for each fatty acid using a 2 way ANOVA (sex x diet) followed by Fisher's LSD test in case of significant interaction.

SFAs: saturated fatty acids; DMAs: dimethyl acetals; MUFAs: monounsaturated fatty acids; PUFAs: polyunsaturated fatty acids.

| Sex | Male | | Female | | 2 way ANOVA | | |
|---|---|---|---|---|---|---|---|
| Diet | n-3 PUFA-sufficient | n-3 PUFA-deficient | n-3 PUFA-sufficient | n-3 PUFA-deficient | Diet effect | Sex effect | Interaction |
| Fatty acid | Mean ± SEM | Mean ± SEM | Mean ± SEM | Mean ± SEM | | | |
| 14:0 | 1.65±0.05[a] | 1.48±0.04[b] | 1.52±0.05[b] | 1.57±0.03[ab] | 0.1824 | 0.668 | *0.0205 |
| 15:0 | 0.08±0.00 | 0.07±0.00 | 0.07±0.00 | 0.06±0.00 | 0.0706 | 0.0706 | 0.5315 |
| 16:0 | 30.14±0.18 | 30.05±0.23 | 29.89±0.30 | 30.01±0.19 | 0.9431 | 0.5409 | 0.6535 |
| 18:0 | 16.08±0.08 | 16.27±0.11 | 0.38±0.02 | 16.20±0.03 | 0.6237 | 0.3995 | 0.1166 |
| **Total SFAs** | **47.94±0.26** | **47.86±0.31** | **47.78±0.34** | **47.84±0.22** | **0.9841** | **0.7559** | **0.8172** |
| DMA 16:0 | 2.10±0.02[b] | 2.19±0.04[a] | 2.13±0.03[ab] | 2.08±0.03[b] | 0.4769 | 0.2032 | *0.0300 |
| DMA 18:0 | 1.42±0.02 | 1.39±0.02 | 1.41±0.01 | 1.36±0.01 | *0.0253 | 0.2106 | 0.6183 |
| DMA 18:1n-9 | 0.34±0.01 | 0.31±0.01 | 0.34±0.01 | 0.33±0.01 | *0.0458 | 0.2812 | 0.2812 |
| DMA 18:1n-7 | 0.37±0.01 | 0.37±0.01 | 0.38±0.02 | 0.39±0.02 | 0.6798 | 0.3213 | 0.518 |
| **Total DMAs** | **4.23±0.02** | **4.26±0.05** | **4.26±0.05** | **4.17±0.02** | **0.4702** | **0.374** | **0.1347** |
| 16:1n-9 | 2.65±0.05 | 2.43±0.06 | 2.54±0.05 | 2.53±0.06 | *0.0494 | 0.9416 | 0.0742 |
| 16:1n-7 | 2.33±0.04 | 2.37±0.04 | 2.28±0.05 | 2.44±0.03 | *0.0249 | 0.7971 | 0.1202 |
| 18:1t | 0.28±0.01 | 0.29±0.01 | 0.30±0.01 | 0.32±0.02 | 0.3917 | 0.1412 | 0.4526 |
| 18:1n-9 | 13.32±0.08 | 12.86±0.12 | 13.36±0.06 | 13.28±0.28 | 0.1121 | 0.1646 | 0.2513 |
| 18:1n-7 | 3.97±0.07 | 3.80±0.05 | 3.98±0.06 | 4.04±0.10 | 0.4527 | 0.104 | 0.1499 |
| 20:1n-9 | 0.28±0.01 | 0.22±0.00 | 0.28±0.01 | 0.23±0.00 | ****<0.0001 | 0.5048 | 0.2357 |
| **Total MUFAs** | **22.83±0.19** | **21.96±0.24** | **22.72±0.19** | **22.84±0.48** | **0.2269** | **0.2114** | **0.1189** |
| 18:2n-6 | 0.38±0.02 | 0.60±0.02 | 0.37±0.01 | 0.64±0.03 | ****<0.0001 | 0.5281 | 0.2604 |

| | | | | | | | |
|---|---|---|---|---|---|---|---|
| 20:3n-6 | 0.13±0.01 | 0.17±0.01 | 0.12±0.01 | 0.17±0.01 | ****<0.0001 | 0.7089 | 0.8517 |
| 20:4n-6 | 10.14±0.09 | 11.30±0.23 | 10.11±0.16 | 10.91±0.21 | ****<0.0001 | 0.2519 | 0.3358 |
| 22:4n-6 | 2.41±0.05 | 2.83±0.08 | 2.51±0.04 | 2.67±0.07 | ***0.0001 | 0.6214 | 0.054 |
| 22:5n-6 | 2.37±0.13 | 5.42±0.20 | 2.46±0.14 | 5.43±0.22 | ****<0.0001 | 0.7636 | 0.8288 |
| **n-6 PUFAs** | **15.52±0.30** | **20.32±0.41** | **15.56±0.19** | **19.82±0.45** | ******<0.0001** | **0.5164** | **0.4616** |
| 18:3n-3 | 0.05±0.01 | 0.04±0.00 | 0.04±0.00 | 0.05±0.01 | 0.6445 | 0.6445 | 0.1013 |
| 20:5n-3 | 0.45±0.01[a] | 0.29±0.01[c] | 0.41±0.02[a] | 0.34±0.03[b] | ****<0.0001 | 0.6432 | *0.0195 |
| 22:5n-3 | 0.11±0.00 | 0.04±0.00 | 0.10±0.00 | 0.04±0.00 | ****<0.0001 | 0.8493 | 0.3474 |
| 22:6n-3 | 8.43±0.21 | 4.95±0.31 | 8.70±0.16 | 4.57±0.24 | ****<0.0001 | 0.814 | 0.1803 |
| **n-3 PUFAs** | **9.04±0.22** | **5.32±0.30** | **9.26±0.18** | **5.00±0.24** | ******<0.0001** | **0.8457** | **0.2647** |
| **n-6/n-3 PUFA** | **1.72±0.06** | **3.89±0.24** | **1.68±0.03** | **4.01±0.22** | ******<0.0001** | **0.812** | **0.6141** |
| 20:3n-9 | 0.46±0.02 | 0.28±0.02 | 0.43±0.02 | 0.34±0.03 | ****<0.0001 | 0.513 | 0.0509 |
| **Total PUFAs** | **25.02±0.30** | **25.92±0.47** | **25.24±0.31** | **25.16±0.46** | **0.3069** | **0.5074** | **0.2249** |

**<u>Supplementary Table 2:</u>** Brain fatty acid composition at E17.5: Data are represented as mean ± SEM (% of total FAMEs and DMAs). N = 6 mice/group.

Statistical comparisons were performed for each fatty acid using a 2 way ANOVA (sex x diet) followed by Fisher's LSD test in case of significant interaction.

SFAs: saturated fatty acids; DMAs: dimethyl acetals; MUFAs: monounsaturated fatty acids; PUFAs: polyunsaturated fatty acids.

| Sex | Male | | Female | | 2 way ANOVA | | |
|---|---|---|---|---|---|---|---|
| Diet | n-3 PUFA-sufficient | n-3 PUFA-deficient | n-3 PUFA-sufficient | n-3 PUFA-deficient | Diet effect | Sex effect | Interaction |
| Fatty acid | Mean ± SEM | Mean ± SEM | Mean ± SEM | Mean ± SEM | | | |
| 14:0 | 0.70±0.08 | 1.77±0.54 | 0.88±0.10 | 2.66±0.13 | ***0.0002 | 0.069 | 0.2148 |
| 15:0 | 0.14±0.01 | 0.20±0.04 | 0.12±0.01 | 0.25±0.05 | **0.0034 | 0.3946 | 0.2043 |
| 16:0 | 31.35±0.28 | 34.60±1.66 | 31.19±0.30 | 37.19±0.15 | ****<0.0001 | 0.166 | 0.1213 |
| 17:0 | 0.14±0.01 | 0.12±0.01 | 0.12±0.02 | 0.12±0.01 | 0.4367 | 0.3526 | 0.5672 |
| 18:0 | 19.41±0.46 | 17.47±0.91 | 18.55±0.68 | 15.97±0.10 | **0.0047 | 0.0994 | 0.6394 |
| 20:0 | 0.16±0.02 | 0.07±0.01 | 0.13±0.03 | 0.14±0.08 | 0.3276 | 0.5816 | 0.1812 |
| Other SFAs | 0.99±0.06 | 2.03±0.57 | 1.13±0.09 | 3.05±0.06 | ***0.0001 | 0.0585 | 0.1411 |
| **Total SFAs** | **51.89±0.45** | **54.22±1.35** | **50.99±0.54** | **56.33±0.02** | *****0.0003** | **0.4507** | **0.0744** |
| DMA 16:0 | 1.43±0.05 | 2.04±0.28 | 1.54±0.09 | 1.58±0.20 | 0.0621 | 0.2907 | 0.0972 |
| DMA 18:0 | 1.22±0.03 | 1.16±0.05 | 1.28±0.06 | 1.05±0.02 | **0.0099 | 0.6444 | 0.1018 |
| DMA 18:1n-9 | 0.45±0.06 | 0.28±0.04 | 0.46±0.03 | 0.22±0.01 | **0.0033 | 0.7475 | 0.4918 |
| DMA 18:1n-7 | 0.38±0.03[c] | 0.61±0.12[b] | 0.44±0.02[bc] | 1.23±0.12[a] | ****<0.0001 | ***0.0005 | **0.0023 |
| **Total DMAs** | **3.48±0.12** | **4.09±0.33** | **3.72±0.13** | **4.08±0.31** | **0.0441** | **0.6041** | **0.5887** |
| 16:1n-9 | 2.77±0.10 | 2.95±0.24 | 2.87±0.06 | 3.43±0.07 | *0.0184 | 0.0552 | 0.185 |
| 16:1n-7 | 2.00±0.10 | 2.64±0.21 | 2.17±0.11 | 3.14±0.11 | ****<0.0001 | *0.0340 | 0.2601 |
| 18:1t | 0.73±0.41 | 0.23±0.00 | 0.25±0.02 | 0.25±0.01 | 0.3362 | 0.3758 | 0.3264 |
| 18:1n-9 | 17.43±0.41 | 15.01±0.42 | 16.84±0.70 | 15.22±0.48 | **0.0033 | 0.7475 | 0.4918 |
| 18:1n-7 | 4.51±0.08 | 4.04±0.13 | 4.51±0.16 | 3.86±0.09 | ****<0.0001 | 0.5035 | 0.4849 |

| | | | | | | | |
|---|---|---|---|---|---|---|---|
| 20:1n-9 | 0.51±0.04 | 0.29±0.01 | 0.45±0.04 | 0.30±0.01 | ***0.0001 | 0.5299 | 0.3501 |
| 20:1n-7 | 0.09±0.01 | 0.08±0.01 | 0.07±0.02 | 0.10±0.01 | 0.4556 | 0.7224 | 0.1465 |
| 22:1n-9 | 0.12±0.01 | 0.08±0.00 | 0.11±0.02 | 0.10±0.02 | 0.0939 | 0.3968 | 0.3249 |
| 24:1n-9 | 4.32±0.36 | 3.70±0.25 | 0.10±0.02 | 0.09±0.03 | 0.3382 | 0.4153 | 0.6273 |
| Other MUFAs | 4.32±0.36 | 3.70±0.25 | 3.86±0.07 | 4.28±0.04 | 0.7036 | 0.7986 | 0.0606 |
| **Total MUFAs** | **28.25±0.44** | **25.39±0.20** | **27.38±0.80** | **26.50±0.30** | ******<0.0001** | **0.8369** | **0.1106** |
| 18:2n-6 | 0.82±0.06 | 1.17±0.03 | 0.79±0.07 | 1.28±0.07 | ****<0.0001 | 0.5863 | 0.283 |
| 20:2n-6 | 0.14±0.03[ab] | 0.10±0.01[b] | 0.09±0.02[b] | 0.22±0.04[a] | 0.1295 | 0.2152 | *0.0116 |
| 20:3n-6 | 0.28±0.02 | 0.29±0.02 | 0.28±0.01 | 0.25±0.01 | 0.5113 | 0.2807 | 0.2807 |
| 20:4n-6 | 6.99±0.29 | 7.27±0.73 | 7.60±0.55 | 5.80±0.23 | 0.1707 | 0.4209 | 0.0684 |
| 22:4n-6 | 1.62±0.05 | 1.74±0.18 | 1.71±0.12 | 1.37±0.07 | 0.3718 | 0.2629 | 0.0819 |
| 22:5n-6 | 1.28±0.09[c] | 3.06±0.37[a] | 1.43±0.10[c] | 2.15±0.13[b] | ****<0.0001 | 0.0772 | *0.0191 |
| **n-6 PUFAs** | **11.14±0.25[b]** | **13.64±1.30[a]** | **11.90±0.61[ab]** | **11.07±0.37[b]** | **0.2871** | **0.2497** | ***0.0450** |
| 20:5n-3 | 0.53±0.02 | 0.32±0.02 | 0.55±0.05 | 0.30±0.02 | ****<0.0001 | 0.9717 | 0.6842 |
| 22:5n-3 | 0.10±0.01 | 0.05±0.01 | 0.12±0.01 | 0.11±0.06 | 0.2071 | 0.0948 | 0.3159 |
| 22:6n-3 | 3.92±0.35 | 2.01±0.26 | 4.63±0.61 | 1.37±0.07 | ****<0.0001 | 0.9415 | 0.1569 |
| **n-3 PUFAs** | **4.55±0.35** | **2.38±0.25** | **5.29±0.57** | **1.78±0.05** | ******<0.0001** | **0.8728** | **0.1447** |
| **n-6/n-3 PUFA** | **2.49±0.13** | **5.76±0.09** | **2.32±0.16** | **6.22±0.18** | ******<0.0001** | **0.3341** | **0.0501** |
| 20:3n-9 | 0.68±0.04 | 0.30±0.03 | 0.72±0.06 | 0.27±0.01 | ****<0.0001 | 0.8405 | 0.502 |
| **Total PUFAs** | **16.37±0.62** | **16.31±1.57** | **17.91±1.13** | **13.12±0.39** | ***0.0457** | **0.4655** | **0.0506** |

**Supplementary Table 3:** Brain fatty acid composition at P0: Data are represented as mean ± SEM (% of total FAMEs and DMAs). N = 3-5 mice/group. Statistical comparisons were performed for each fatty acid using a 2 way ANOVA (sex x diet) followed by Fisher's LSD test in case of significant interaction. SFAs: saturated fatty acids; DMAs: dimethyl acetals; MUFAs: monounsaturated fatty acids; PUFAs: polyunsaturated fatty acids.

| | Sex | Male | | Female | | 2 way ANOVA | | |
|---|---|---|---|---|---|---|---|---|
| | Diet | n-3 PUFA-sufficient | n-3 PUFA-deficient | n-3 PUFA-sufficient | n-3 PUFA-deficient | Diet effect | Sex effect | Interaction |
| | Oxylipin | Mean ± SEM | Mean ± SEM | Mean ± SEM | Mean ± SEM | | | |
| LA | 9-HODE | 936±100 | 2192±176 | 1331±179 | 2910±472 | ****<0.0001 | *0.025 | 0.4758 |
| | 13-HODE | 976±88 | 2141±175 | 1285±185 | 3568±1019 | ***0.0005 | *0.0376 | 0.1597 |
| AA | 5,6-EET | 185±7 | 226±24 | 226±29 | 247±17 | 0.184 | 0.1942 | 0.6528 |
| | 8,9-EET | 356±16 | 496±58 | 506±109 | 655±47 | 0.0762 | 0.0603 | 0.9476 |
| | 11,12-EET | 248±31 | 342±6 | 346±68 | 451±32 | 0.054 | *0.0468 | 0.906 |
| | 14,15-EET | 175±15 | 231±13 | 194±34 | 271±43 | *0.0300 | 0.3053 | 0.7126 |
| | 5-HETE | 4852±363[b] | 8804±1193[a] | 7657±716[ab] | 6987±318[a] | 0.0502 | 0.5278 | **0.0095 |
| | 8-HETE | 769±80 | 898±145 | 688±80 | 791±64 | *0.0156 | 0.6293 | 0.1216 |
| | 12-HETE | 3969±451 | 5444±1040 | 5420±615 | 5801±796 | 0.2275 | 0.2391 | 0.4688 |
| | 15-HETE | 5022±635 | 9352±1406 | 7420±712 | 7883±784 | *0.0228 | 0.6255 | 0.0577 |
| | 5oxoETE | 3963±322 | 5320±762 | 4617±554 | 6131±868 | *0.0351 | 0.2516 | 0.8997 |
| | LxA4 | 363±44 | 575±81 | 451±72 | 622±64 | *0.0148 | 0.3368 | 0.7666 |
| | LXB4 | 412±44 | 425±27 | 463±62 | 470±149 | 0.8866 | 0.5046 | 0.9738 |
| | 8isoPGA2 | 404±45 | 591±80 | 599±92 | 471±91 | 0.7207 | 0.648 | 0.0701 |
| | PGD2 | 1397±262 | 1705±350 | 1346±290 | 2792±779 | *0.0438 | 0.21 | 0.1712 |
| | PGE2 | 465±84 | 500±67 | 480±98 | 835±135 | 0.0656 | 0.0947 | 0.1228 |
| | PGF2a | 1176±174 | 1170±144 | 1342±238 | 1632±224 | 0.5078 | 0.1551 | 0.4902 |

| | | | | | | | | |
|---|---|---|---|---|---|---|---|---|
| | 11B-PGF2a | 47±32 | 29±29 | 74±20 | 23±23 | 0.2393 | 0.6975 | 0.5718 |
| | 15dPGJ2 | 45±7 | 65±6 | 69±11 | 51±11 | 0.9394 | 0.5734 | 0.0596 |
| | 6kPGF1a | 786±159 | 541±17 | 1107±273 | 889±167 | 0.2747 | 0.1229 | 0.9502 |
| | TXB2 | 1164±104 | 1139±164 | 1343±199 | 1374±321 | 0.9888 | 0.2995 | 0.8873 |
| EPA | 18-HEPE | 31±13 | 0±0 | 55±25 | 0±0 | *0.0291 | 0.516 | 0.516 |
| DHA | 14-HDoHE | 769±80 | 636±98 | 1057±150 | 512±39 | *0.0113 | 0.4866 | 0.0967 |
| | 17-HDoHE | 649±123 | 446±81 | 864±102 | 390±85 | **0.0086 | 0.4795 | 0.2366 |

**Supplementary Table 4:** Brain oxylipin profile at P0: Data are represented as mean ± SEM (pg/mg of protein). N = 3-5 mice/group. Statistical comparisons were performed for each oxylipin using a 2 way ANOVA (sex x diet) followed by Fisher's LSD test in case of significant interaction. LA: linoleic acid; AA: arachidonic acid; EPA: eicosapentaenoic acid; DHA: docosahexaenoic acid.

| Sex | Male | | Female | | 2 way ANOVA | | |
|---|---|---|---|---|---|---|---|
| Diet | n-3 PUFA-sufficient | n-3 PUFA-deficient | n-3 PUFA-sufficient | n-3 PUFA-deficient | Diet effect | Sex effect | Interaction |
| Fatty acid | Mean ± SEM | Mean ± SEM | Mean ± SEM | Mean ± SEM | | | |
| 14:0 | 1.76±0.10 | 2.30±0.17 | 1.76±0.11 | 2.08±0.17 | **0.0082 | 0.4647 | 0.4325 |
| 15:0 | 0.11±0.01 | 0.13±0.01 | 0.11±0.00 | 0.12±0.01 | 0.2361 | 0.78 | 0.78 |
| 16:0 | 31.72±0.15 | 32.27±0.33 | 31.65±0.17 | 33.42±0.98 | *0.0433 | 0.3255 | 0.2701 |
| 17:0 | 0.10±0.01 | 0.10±0.01 | 0.10±0.01 | 0.10±0.01 | 0.7147 | 0.9028 | 0.7147 |
| 18:0 | 16.74±0.18 | 16.70±0.10 | 16.88±0.22 | 16.89±0.41 | 0.9693 | 0.5359 | 0.9265 |
| 20:0 | 0.08±0.03 | 0.07±0.02 | 0.11±0.03 | 0.11±0.04 | 0.8978 | 0.3421 | 0.8978 |
| Other SFAs | 1.95±0.09 | 2.50±0.19 | 1.98±0.09 | 2.31±0.18 | **0.0077 | 0.5875 | 0.4417 |
| **Total SFAs** | **50.50±0.21** | **51.58±0.53** | **50.61±0.31** | **52.71±1.53** | **0.074** | **0.4679** | **0.5464** |
| DMA 16:0 | 1.92±0.09 | 1.91±0.08 | 2.00±0.05 | 2.18±0.05 | 0.2731 | *0.0291 | 0.1929 |
| DMA 18:0 | 1.45±0.06 | 1.39±0.08 | 1.49±0.04 | 1.57±0.04 | 0.8399 | 0.078 | 0.2234 |
| DMA 18:1n-9 | 0.26±0.02 | 0.29±0.03 | 0.22±0.02 | 0.30±0.03 | *0.0381 | 0.6919 | 0.2751 |
| DMA 18:1n-7 | 0.25±0.01 | 0.30±0.04 | 0.23±0.01 | 0.29±0.04 | 0.0774 | 0.6434 | 0.8423 |
| **Total DMAs** | **3.88±0.07** | **3.88±0.18** | **3.93±0.07** | **4.34±0.11** | **0.0959** | ***0.0413** | **0.0959** |
| 16:1n-9 | 2.47±0.09 | 2.55±0.06 | 2.41±0.09 | 2.63±0.11 | 0.1304 | 0.9486 | 0.4309 |
| 16:1n-7 | 1.91±0.11 | 2.46±0.05 | 1.81±0.11 | 2.49±0.09 | ****<0.0001 | 0.6966 | 0.4905 |
| 18:1t | 0.15±0.01 | 0.17±0.02 | 0.14±0.01 | 0.20±0.03 | *0.0497 | 0.4894 | 0.3685 |
| 18:1n-9 | 11.62±0.09 | 10.52±0.10 | 11.71±0.14 | 11.01±0.18 | ****<0.0001 | *0.0439 | 0.1555 |
| 18:1n-7 | 3.28±0.04 | 3.11±0.05 | 3.28±0.06 | 3.16±0.09 | *0.0402 | 0.7053 | 0.7053 |

| | | | | | | | |
|---|---|---|---|---|---|---|---|
| 20:1n-9 | 0.26±0.01 | 0.21±0.03 | 0.28±0.00 | 0.25±0.04 | 0.126 | 0.37 | 0.7625 |
| 20:1n-7 | 0.03±0.00 | 0.03±0.01 | 0.03±0.00 | 0.03±0.00 | 0.5636 | 0.5636 | 0.5636 |
| 22:1n-9 | 0.05±0.00 | 0.05±0.00 | 0.05±0.00 | 0.05±0.00 | 0.1598 | 0.7719 | 0.3894 |
| 24:1n-9 | 0.04±0.00 | 0.03±0.01 | 0.04±0.00 | 0.03±0.01 | 0.1984 | 0.3844 | 0.6607 |
| Other MUFAs | 3.00±0.09 | 3.04±0.10 | 2.96±0.09 | 3.18±0.18 | 0.3001 | 0.6859 | 0.4596 |
| **Total MUFAs** | **19.81±0.24** | **19.12±0.16** | **19.76±0.32** | **19.84±0.51** | **0.3842** | **0.334** | **0.2713** |
| 18:2n-6 | 0.59±0.02 | 0.85±0.02 | 0.60±0.02 | 0.83±0.01 | ****<0.0001 | 0.7155 | 0.3289 |
| 20:2n-6 | 0.11±0.01[ab] | 0.07±0.00[b] | 0.10±0.02[ab] | 0.13±0.01[a] | 0.7068 | 0.0736 | *0.0304 |
| 20:3n-6 | 0.42±0.02 | 0.39±0.01 | 0.43±0.02 | 0.41±0.01 | 0.1477 | 0.5441 | 0.8679 |
| 20:4n-6 | 11.43±0.13 | 12.14±0.20 | 11.47±0.10 | 12.05±0.13 | ***0.0005 | 0.8607 | 0.6667 |
| 22:4n-6 | 2.34±0.04 | 2.88±0.07 | 2.32±0.06 | 2.97±0.07 | ****<0.0001 | 0.6005 | 0.3806 |
| 22:5n-6 | 1.45±0.06 | 5.05±0.24 | 1.45±0.07 | 4.85±0.42 | ****<0.0001 | 0.6885 | 0.6885 |
| **n-6 PUFAs** | **16.35±0.17** | **21.39±0.17** | **16.38±0.21** | **18.99±1.85** | *****0.0009** | **0.2261** | **0.215** |
| 20:5n-3 | 0.31±0.01 | 0.17±0.01 | 0.32±0.03 | 0.26±0.03 | ***0.0004 | 0.0669 | 0.1078 |
| 22:5n-3 | 0.17±0.00 | 0.04±0.00 | 0.17±0.01 | 0.04±0.00 | ****<0.0001 | 0.7719 | 0.3894 |
| 22:6n-3 | 8.45±0.24 | 3.55±0.39 | 8.33±0.24 | 3.57±0.36 | ****<0.0001 | 0.8708 | 0.8219 |
| **n-3 PUFAs** | **8.94±0.22** | **3.76±0.40** | **8.82±0.25** | **3.86±0.37** | ******<0.0001** | **0.9803** | **0.7395** |
| **n-6/n-3 PUFA** | **1.83±0.06** | **5.97±0.69** | **1.86±0.07** | **5.07±0.73** | ******<0.0001** | **0.3967** | **0.366** |
| 20:3n-9 | 0.52±0.03 | 0.27±0.02 | 0.50±0.04 | 0.25±0.01 | ****<0.0001 | 0.4948 | >0.9999 |
| **Total PUFAs** | **25.81±0.10** | **25.43±0.40** | **25.70±0.14** | **23.11±1.99** | **0.1634** | **0.2515** | **0.2937** |

**<u>Supplementary Table 5:</u>** Brain fatty acid composition at P7: Data are represented as mean ± SEM (% of total FAMEs and DMAs). N = 5 mice/group. Statistical comparisons were performed for each fatty acid using a 2 way ANOVA (sex x diet) followed by Fisher's LSD test in case of significant interaction. SFAs: saturated fatty acids; DMAs: dimethyl acetals; MUFAs: monounsaturated fatty acids; PUFAs: polyunsaturated fatty acids.

| | Sex | Male | | Female | | 2 way ANOVA | | |
|---|---|---|---|---|---|---|---|---|
| | Diet | n-3 PUFA-sufficient | n-3 PUFA-deficient | n-3 PUFA-sufficient | n-3 PUFA-deficient | Diet effect | Sex effect | Interaction |
| | Oxylipin | Mean ± SEM | Mean ± SEM | Mean ± SEM | Mean ± SEM | | | |
| LA | 9-HODE | 1035±79 | 2208±149 | 794±85 | 2285±387 | ****<0.0001 | 0.7089 | 0.4729 |
| | 13-HODE | 916±76 | 2023±200 | 651±62 | 1907±367 | ****<0.0001 | 0.3885 | 0.7333 |
| AA | 5,6-EET | 479±67 | 409±75 | 366±76 | 535±42 | 0.4673 | 0.9183 | 0.0911 |
| | 8,9-EET | 504±43 | 509±34 | 430±92 | 661±100 | 0.1248 | 0.6032 | 0.1412 |
| | 11,12-EET | 183±21 | 237±36 | 153±16 | 225±56 | 0.0963 | 0.5635 | 0.8087 |
| | 14,15-EET | 111±14 | 125±22 | 95±11 | 125±35 | 0.3367 | 0.7279 | 0.7314 |
| | 5-HETE | 12654±1869 | 19828±3111 | 8544±849 | 17520±5039 | *0.0203 | 0.321 | 0.7774 |
| | 8-HETE | 1127±206 | 1562±155 | 712±73 | 1540±410 | *0.0202 | 0.3863 | 0.435 |
| | 12-HETE | 4768±816 | 5635±326 | 2845±290 | 6255±1363 | *0.0195 | 0.4405 | 0.1423 |
| | 15-HETE | 11785±1600 | 16477±1240 | 7881±906 | 16034±3733 | **0.0092 | 0.3316 | 0.4369 |
| | 5oxoETE | 8356±1121 | 8722±534 | 7823±925 | 8451±726 | 0.5689 | 0.6445 | 0.8803 |
| | LxA4 | 676±117 | 1077±178 | 534±48 | 811±244 | 0.0551 | 0.2319 | 0.7102 |
| | LXB4 | 675±97[ab] | 498±117[b] | 564±96[b] | 886±113[a] | 0.5036 | 0.2086 | *0.0315 |
| | 8isoPGA2 | 561±125 | 909±141 | 481±62 | 851±203 | *0.0223 | 0.636 | 0.9394 |
| | PGD2 | 2827±611 | 3508±414 | 2716±321 | 2866±470 | 0.3858 | 0.4313 | 0.5764 |
| | PGE2 | 851±97 | 1203±209 | 797±75 | 892±203 | 0.1768 | 0.2655 | 0.4263 |
| | PGF2a | 2516±480 | 1782±166 | 2263±300 | 2216±297 | 0.2543 | 0.7887 | 0.3139 |

| | | | | | | | | |
|---|---|---|---|---|---|---|---|---|
| | 11B-PGF2a | 5±5 | 0±0 | 0±0 | 12±12 | 0.6362 | 0.6362 | 0.2023 |
| | 15dPGJ2 | 54±7 | 52±5 | 43±8 | 79±15 | 0.0956 | 0.4086 | 0.0562 |
| | 6kPGF1a | 548±80 | 632±73 | 479±38 | 724±80 | *0.0328 | 0.8674 | 0.2681 |
| | TXB2 | 1812±278 | 1860±233 | 1586±197 | 1970±365 | 0.445 | 0.8368 | 0.5496 |
| EPA | 18-HEPE | 41±28 | 0±0 | 34±21 | 0±0 | *0.0467 | 0.8524 | 0.8524 |
| DHA | 14-HDoHE | 1315±234[a] | 568±46[b] | 766±74[b] | 596±101[b] | **0.0036 | 0.0708 | *0.0477 |
| | 17-HDoHE | 769±128[a] | 320±47[b] | 456±100[b] | 437±93[b] | *0.0273 | 0.3266 | *0.0404 |

**Supplementary Table 6:** Brain oxylipin profile at P7: Data are represented as mean ± SEM (pg/mg of protein). N = 5 mice/group. Statistical comparisons were performed for each oxylipin using a 2 way ANOVA (sex x diet) followed by Fisher's LSD test in case of significant interaction. LA: linoleic acid; AA: arachidonic acid; EPA: eicosapentaenoic acid; DHA: docosahexaenoic acid.

| Sex | Male | | Female | | 2 way ANOVA | | |
|---|---|---|---|---|---|---|---|
| **Diet** | **n-3 PUFA-sufficient** | **n-3 PUFA-deficient** | **n-3 PUFA-sufficient** | **n-3 PUFA-deficient** | **Diet effect** | **Sex effect** | **Interaction** |
| **Fatty acid** | **Mean ± SEM** | **Mean ± SEM** | **Mean ± SEM** | **Mean ± SEM** | | | |
| 14:0 | 0.42±0.04 | 0.61±0.06 | 0.46±0.05 | 0.60±0.05 | **0.0034 | 0.7448 | 0.6261 |
| 15:0 | 0.08±0.01 | 0.08±0.01 | 0.08±0.01 | 0.09±0.01 | 0.5885 | 0.4198 | 0.7861 |
| 16:0 | 26.37±0.66 | 26.92±0.38 | 25.26±0.47 | 26.83±0.19 | *0.0338 | 0.2062 | 0.2828 |
| 17:0 | 0.12±0.01 | 0.12±0.00 | 0.12±0.01 | 0.12±0.01 | 0.7979 | 0.7979 | 0.7979 |
| 18:0 | 19.90±0.35 | 19.11±0.19 | 19.34±0.22 | 19.14±0.15 | 0.0575 | 0.2825 | 0.2372 |
| 20:0 | 0.15±0.01 | 0.16±0.05 | 0.13±0.01 | 0.17±0.05 | 0.4804 | 0.932 | 0.712 |
| Other SFAs | 0.09±0.01 | 0.06±0.00 | 0.07±0.01 | 0.06±0.01 | *0.0179 | 0.3071 | 0.2058 |
| **Total SFAs** | **0.86±0.04** | **1.04±0.11** | **0.87±0.03** | **1.05±0.04** | ***0.0121** | **0.8381** | **0.9624** |
| DMA 16:0 | 47.13±0.96 | 47.07±0.60 | 45.47±0.31 | 47.02±0.28 | 0.2354 | 0.1731 | 0.1981 |
| DMA 18:0 | 2.35±0.06 | 2.47±0.04 | 2.41±0.02 | 2.55±0.05 | *0.0139 | 0.1615 | 0.8986 |
| DMA 18:1n-9 | 2.37±0.12 | 2.19±0.04 | 2.49±0.11 | 2.13±0.02 | **0.0054 | 0.6868 | 0.2829 |
| DMA 18:1n-7 | 0.67±0.03 | 0.53±0.01 | 0.70±0.03 | 0.51±0.04 | ****<0.0001 | 0.8922 | 0.421 |
| **Total DMAs** | **0.49±0.02** | **0.44±0.01** | **0.53±0.01** | **0.43±0.04** | ****0.0036** | **0.6411** | **0.3778** |
| 16:1n-9 | 5.88±0.20 | 5.63±0.07 | 6.13±0.13 | 5.62±0.09 | *0.0104 | 0.3811 | 0.3282 |
| 16:1n-7 | 0.89±0.07 | 1.14±0.03 | 0.92±0.10 | 1.08±0.02 | **0.0040 | 0.7993 | 0.4873 |
| 18:1t | 0.65±0.04 | 0.83±0.03 | 0.69±0.06 | 0.82±0.02 | **0.0012 | 0.7831 | 0.5018 |
| 18:1n-9 | 0.09±0.01 | 0.20±0.06 | 0.09±0.00 | 0.14±0.02 | *0.0158 | 0.3382 | 0.308 |
| 18:1n-7 | 12.73±0.39 | 10.71±0.16 | 11.78±0.07 | 10.39±0.08 | ****<0.0001 | *0.0106 | 0.1691 |

| | | | | | | | |
|---|---|---|---|---|---|---|---|
| 20:1n-9 | 3.02±0.03 | 3.00±0.05 | 2.92±0.02 | 2.91±0.03 | 0.6607 | *0.0163 | 0.8259 |
| 20:1n-7 | 0.38±0.03 | 0.34±0.08 | 0.39±0.03 | 0.33±0.10 | 0.4543 | 0.9646 | 0.894 |
| 22:1n-9 | 0.08±0.01 | 0.05±0.01 | 0.07±0.01 | 0.06±0.01 | *0.0293 | 0.9013 | 0.7104 |
| 24:1n-9 | 0.07±0.00 | 0.05±0.01 | 0.07±0.00 | 0.05±0.00 | **0.0012 | 0.7962 | 0.7962 |
| Other MUFAs | 1.50±0.06 | 1.79±0.13 | 1.54±0.07 | 1.66±0.10 | *0.0414 | 0.648 | 0.3938 |
| **Total MUFAs** | **17.91±0.43** | **1.79±0.13** | **16.93±0.18** | **15.78±0.13** | *****0.0002** | ***0.0155** | **0.4551** |
| 18:2n-6 | 0.86±0.03[b] | 0.98±0.03[a] | 0.74±0.01[c] | 1.02±0.01[a] | ****<0.0001 | 0.0722 | **0.0025 |
| 20:2n-6 | 0.16±0.03 | 0.18±0.05 | 0.16±0.04 | 0.18±0.02 | 0.6028 | 0.9537 | 0.9537 |
| 20:3n-6 | 0.60±0.01 | 0.48±0.02 | 0.56±0.02 | 0.48±0.01 | ****<0.0001 | 0.1751 | 0.1397 |
| 20:4n-6 | 12.36±0.32 | 13.84±0.33 | 13.43±0.11 | 14.06±0.24 | **0.0010 | *0.0278 | 0.1259 |
| 22:4n-6 | 2.55±0.04 | 3.56±0.09 | 2.69±0.06 | 3.62±0.06 | ****<0.0001 | 0.1403 | 0.4992 |
| 22:5n-6 | 1.56±0.17 | 8.27±0.53 | 1.75±0.11 | 8.42±0.32 | ****<0.0001 | 0.611 | 0.9494 |
| **n-6 PUFAs** | **18.10±0.47** | **27.32±0.87** | **19.33±0.18** | **27.78±0.46** | ******<0.0001** | **0.1467** | **0.4942** |
| 20:5n-3 | 0.28±0.01[a] | 0.20±0.01[b] | 0.30±0.01[a] | 0.18±0.01[b] | ****<0.0001 | 0.9036 | *0.0199 |
| 22:5n-3 | 0.14±0.00 | 0.03±0.00 | 0.15±0.01 | 0.04±0.00 | ****<0.0001 | 0.0599 | 0.3709 |
| 22:6n-3 | 10.24±0.55 | 3.26±0.10 | 11.31±0.28 | 3.44±0.19 | ****<0.0001 | 0.0731 | 0.1909 |
| **n-3 PUFAs** | **10.66±0.55** | **3.49±0.10** | **11.76±0.26** | **3.65±0.19** | ******<0.0001** | **0.0671** | **0.1657** |
| **n-6/n-3 PUFA** | **1.71±0.05** | **7.88±0.45** | **1.65±0.03** | **7.71±0.50** | ******<0.0001** | **0.7266** | **0.8656** |
| 20:3n-9 | 0.31±0.01[b] | 0.17±0.01[c] | 0.38±0.04[a] | 0.15±0.01[c] | ****<0.0001 | 0.2169 | *0.0449 |
| **Total PUFAs** | **29.07±0.99** | **30.98±0.80** | **31.48±0.40** | **31.58±0.37** | **0.1645** | ***0.0443** | **0.2099** |

**Supplementary Table 7:** Hippocampal fatty acid composition at P14: Data are represented as mean ± SEM (% of total FAMEs and DMAs). N = 5 mice/group.

Statistical comparisons were performed for each fatty acid using a 2 way ANOVA (sex x diet) followed by Fisher's LSD test in case of significant interaction.

SFAs: saturated fatty acids; DMAs: dimethyl acetals; MUFAs: monounsaturated fatty acids; PUFAs: polyunsaturated fatty acids.

| | Sex | Male | | Female | | 2 way ANOVA | | |
|---|---|---|---|---|---|---|---|---|
| | Diet | n-3 PUFA-sufficient | n-3 PUFA-deficient | n-3 PUFA-sufficient | n-3 PUFA-deficient | Diet effect | Sex effect | Interaction |
| | Oxylipin | Mean ± SEM | Mean ± SEM | Mean ± SEM | Mean ± SEM | | | |
| LA | 9-HODE | 2315±163 | 3202±810 | 2360±242 | 3397±408 | 0.0605 | 0.8046 | 0.8765 |
| | 13-HODE | 2549±303 | 3038±660 | 2416±236 | 3289±475 | 0.1494 | 0.8967 | 0.6746 |
| AA | 5,6-EET | 880±77 | 1078±285 | 711±81 | 1197±194 | 0.0773 | 0.8919 | 0.4394 |
| | 8,9-EET | 2309±336 | 2360±622 | 1867±247 | 2516±433 | 0.4305 | 0.7455 | 0.4998 |
| | 11,12-EET | 1385±180 | 1226±332 | 1016±112 | 1428±233 | 0.5888 | 0.7217 | 0.2306 |
| | 14,15-EET | 664±61 | 670±204 | 603±84 | 749±131 | 0.5747 | 0.9462 | 0.6014 |
| | 5-HETE | 18403±1864 | 22905±7534 | 20011±2942 | 23842±3685 | 0.3724 | 0.7829 | 0.942 |
| | 8-HETE | 1374±150 | 1522±456 | 1458±262 | 1525±261 | 0.7268 | 0.8878 | 0.8953 |
| | 12-HETE | 9352±1875 | 7422±1543 | 6807±1102 | 9191±1815 | 0.89 | 0.813 | 0.1998 |
| | 15-HETE | 13098±1145 | 14078±4114 | 13543±2198 | 14530±2344 | 0.7178 | 0.8688 | 0.9989 |
| | 5oxoETE | 17312±1914 | 14604±3946 | 14518±2214 | 15161±2673 | 0.7169 | 0.6945 | 0.5575 |
| | LxA4 | 942±143 | 1021±197 | 1018±169 | 1073±309 | 0.7586 | 0.7689 | 0.9576 |
| | LXB4 | 2604±361 | 2982±695 | 2288±446 | 3125±682 | 0.2983 | 0.8806 | 0.6902 |
| | 8isoPGA2 | 2207±319 | 2023±469 | 1780±217 | 2023±325 | 0.9323 | 0.5449 | 0.5451 |
| | PGD2 | 3276±728 | 3155±1150 | 2531±570 | 2030±647 | 0.7049 | 0.2625 | 0.8161 |
| | PGE2 | 1853±365 | 1708±544 | 1256±235 | 1565±557 | 0.8566 | 0.4191 | 0.6168 |
| | PGF2a | 17150±2284 | 14705±3470 | 13475±3080 | 16685±4208 | 0.9102 | 0.8025 | 0.4088 |

| | | | | | | | | |
|---|---|---|---|---|---|---|---|---|
| | 15dPGJ2 | 205±29 | 167±37 | 168±21 | 161±25 | 0.439 | 0.4558 | 0.5826 |
| | 6kPGF1a | 3717±1402 | 1631±675 | 3779±1189 | 2040±536 | 0.0779 | 0.8199 | 0.8663 |
| | TXB2 | 5528±670 | 6033±1434 | 5026±513 | 5683±957 | 0.5534 | 0.6633 | 0.9379 |
| DHA | 14-HDoHE | 1660±261 | 489±147 | 1744±350 | 514±99 | ***0.0001 | 0.8213 | 0.9016 |
| | 17-HDoHE | 1419±145 | 408±117 | 1654±387 | 486±87 | ***0.0001 | 0.4847 | 0.7253 |

**Supplementary Table 8:** Hippocampal oxylipin profile at P14: Data are represented as mean ± SEM (pg/mg of protein). N = 5 mice/group. Statistical comparisons were performed for each oxylipin using a 2 way ANOVA (sex x diet). LA: linoleic acid; AA: arachidonic acid; DHA: docosahexaenoic acid.

| Sex | Male | | Female | | 2 way ANOVA | | |
|---|---|---|---|---|---|---|---|
| Diet | n-3 PUFA-sufficient | n-3 PUFA-deficient | n-3 PUFA-sufficient | n-3 PUFA-deficient | Diet effect | Sex effect | Interaction |
| Fatty acid | Mean ± SEM | Mean ± SEM | Mean ± SEM | Mean ± SEM | | | |
| 14:0 | 0.22±0.02[b] | 0.28±0.02[a] | 0.26±0.01[ab] | 0.26±0.02[b] | 0.4529 | 0.4529 | **0.0050 |
| 15:0 | 0.06±0.00[b] | 0.07±0.011[a] | 0.07±0.00[ab] | 0.06±0.00[ab] | 0.6126 | >0.9999 | *0.0201 |
| 16:0 | 23.21±0.17 | 24.62±0.13 | 23.42±0.22 | 24.15±0.21 | ****<0.0001 | 0.4981 | 0.0825 |
| 17:0 | 0.13±0.00 | 0.14±0.01 | 0.14±0.00 | 0.14±0.01 | 0.8392 | 0.5447 | 0.3177 |
| 18:0 | 19.72±0.15 | 19.62±0.09 | 19.76±0.15 | 19.83±0.07 | 0.9202 | 0.3164 | 0.4566 |
| 20:0 | 0.29±0.01 | 0.32±0.04 | 0.41±0.03 | 0.41±0.04 | 0.5723 | **0.0051 | 0.6551 |
| Other SFAs | 0.17±0.01 | 0.16±0.01 | 0.18±0.02 | 0.16±0.01 | 0.1416 | 0.5018 | 0.8657 |
| **Total SFAs** | **0.87±0.01** | **0.98±0.05** | **1.06±0.04** | **1.04±0.06** | **0.4059** | ***0.0134** | **0.174** |
| DMA 16:0 | 43.80±0.29 | 45.22±0.17 | 44.24±0.35 | 45.03±0.16 | ***0.0005 | 0.6416 | 0.2317 |
| DMA 18:0 | 2.61±0.03 | 2.76±0.04 | 2.65±0.04 | 2.71±0.04 | *0.0135 | >0.9999 | 0.2373 |
| DMA 18:1n-9 | 3.04±0.05 | 2.74±0.06 | 2.87±0.09 | 2.82±0.07 | *0.0227 | 0.5033 | 0.0921 |
| DMA 18:1n-7 | 0.91±0.03 | 0.81±0.02 | 0.97±0.03 | 0.81±0.03 | ***0.0003 | 0.2384 | 0.3588 |
| **Total DMAs** | **0.63±0.02** | **0.60±0.02** | **0.64±0.02** | **0.59±0.02** | **0.0579** | **0.9199** | **0.6883** |
| 16:1n-9 | 7.18±0.07 | 6.90±0.07 | 7.13±0.15 | 6.93±0.10 | *0.0366 | 0.9255 | 0.695 |
| 16:1n-7 | 0.41±0.02 | 0.46±0.02 | 0.45±0.02 | 0.42±0.02 | 0.5683 | 0.7945 | 0.0555 |
| 18:1t | 0.49±0.01[a] | 0.57±0.02[a] | 0.52±0.02[a] | 0.49±0.02[a] | 0.2268 | 0.1921 | *0.0186 |
| 18:1n-9 | 0.11±0.01 | 0.12±0.02 | 0.12±0.01 | 0.15±0.01 | 0.2671 | 0.2671 | 0.5151 |
| 18:1n-7 | 12.85±0.10 | 11.61±0.13 | 13.13±0.23 | 11.82±0.12 | ****<0.0001 | 0.1289 | 0.8205 |

| | | | | | | | |
|---|---|---|---|---|---|---|---|
| 20:1n-9 | 3.21±0.03 | 3.48±0.04 | 3.26±0.03 | 3.46±0.05 | ****<0.0001 | 0.5921 | 0.3143 |
| 20:1n-7 | 0.61±0.03 | 0.48±0.02 | 0.63±0.05 | 0.50±0.05 | **0.0040 | 0.5385 | 0.9589 |
| 22:1n-9 | 0.18±0.01 | 0.16±0.01 | 0.19±0.02 | 0.17±0.02 | 0.1457 | 0.5493 | 0.8804 |
| 24:1n-9 | 0.13±0.01 | 0.12±0.00 | 0.14±0.01 | 0.12±0.01 | 0.202 | 0.4364 | 0.6019 |
| Other MUFAs | 1.44±0.04 | 1.34±0.04 | 1.52±0.06 | 1.36±0.08 | *0.0405 | 0.3785 | 0.585 |
| **Total MUFAs** | **17.99±0.13** | **16.99±0.18** | **18.43±0.29** | **17.13±0.22** | ******<0.0001** | **0.1943** | **0.4779** |
| 18:2n-6 | 0.61±0.01 | 0.81±0.03 | 0.61±0.04 | 0.78±0.02 | ****<0.0001 | 0.7419 | 0.6929 |
| 20:2n-6 | 0.13±0.02 | 0.16±0.01 | 0.13±0.02 | 0.16±0.02 | 0.1117 | >0.9999 | 0.912 |
| 20:3n-6 | 0.51±0.01 | 0.38±0.01 | 0.52±0.01 | 0.39±0.01 | ****<0.0001 | 0.2671 | 0.918 |
| 20:4n-6 | 12.41±0.09 | 12.46±0.26 | 12.20±0.23 | 12.65±0.30 | 0.3048 | 0.9667 | 0.4089 |
| 22:4n-6 | 2.62±0.02 | 3.50±0.06 | 2.60±0.03 | 3.52±0.02 | ****<0.0001 | 0.978 | 0.6796 |
| 22:5n-6 | 1.57±0.06 | 9.64±0.38 | 1.56±0.06 | 9.42±0.14 | ****<0.0001 | 0.5892 | 0.6483 |
| **n-6 PUFAs** | **17.85±0.11** | **26.95±0.47** | **17.62±0.32** | **26.93±0.30** | ******<0.0001** | **0.7116** | **0.7567** |
| 20:5n-3 | 0.34±0.01 | 0.25±0.01 | 0.35±0.03 | 0.26±0.02 | ***0.0002 | 0.409 | 0.8675 |
| 22:5n-3 | 0.13±0.00 | 0.03±0.00 | 0.13±0.00 | 0.03±0.00 | ****<0.0001 | 0.5256 | 0.5256 |
| 22:6n-3 | 12.43±0.24 | 3.53±0.27 | 11.78±0.20 | 3.56±0.30 | ****<0.0001 | 0.2438 | 0.2003 |
| **n-3 PUFAs** | **12.90±0.24** | **3.81±0.28** | **12.26±0.20** | **3.86±0.30** | ******<0.0001** | **0.2691** | **0.1993** |
| **n-6/n-3 PUFA** | **1.39±0.02** | **7.25±0.59** | **1.44±0.02** | **7.15±0.57** | ******<0.0001** | **0.9578** | **0.8589** |
| 20:3n-9 | 0.28±0.01 | 0.11±0.01 | 0.30±0.01 | 0.13±0.00 | ****<0.0001 | 0.0898 | 0.7227 |
| **Total PUFAs** | **31.03±0.32** | **30.88±0.26** | **30.18±0.47** | **30.92±0.33** | **0.4147** | **0.2695** | **0.2246** |

**<u>Supplementary Table 9:</u>** Hippocampal fatty acid composition at P21: Data are represented as mean ± SEM (% of total FAMEs and DMAs). N = 5 mice/group.

Statistical comparisons were performed for each fatty acid using a 2 way ANOVA (sex x diet) followed by Fisher's LSD test in case of significant interaction.

SFAs: saturated fatty acids; DMAs: dimethyl acetals; MUFAs: monounsaturated fatty acids; PUFAs: polyunsaturated fatty acids.

| | Sex | Male | | Female | | 2 way ANOVA | | |
|---|---|---|---|---|---|---|---|---|
| | Diet | n-3 PUFA-sufficient | n-3 PUFA-deficient | n-3 PUFA-sufficient | n-3 PUFA-deficient | Diet effect | Sex effect | Interaction |
| | Oxylipin | Mean ± SEM | Mean ± SEM | Mean ± SEM | Mean ± SEM | | | |
| LA | 9-HODE | 2296±282 | 2184±385 | 1772±140 | 2258±258 | 0.4975 | 0.416 | 0.2829 |
| | 13-HODE | 2124±269 | 2051±448 | 1694±91 | 2076±195 | 0.5627 | 0.448 | 0.396 |
| AA | 5,6-EET | 882±94 | 1009±165 | 814±73 | 1080±444 | 0.4511 | 0.9958 | 0.7862 |
| | 8,9-EET | 2241±298 | 2778±541 | 2560±363 | 3472±1613 | 0.4443 | 0.5908 | 0.8416 |
| | 11,12-EET | 996±189 | 1553±336 | 1207±134 | 2286±1131 | 0.214 | 0.4658 | 0.6846 |
| | 14,15-EET | 570±75 | 841±180 | 702±76 | 1248±638 | 0.2642 | 0.4563 | 0.7013 |
| | 5-HETE | 22059±2194 | 16802±2940 | 16871±1544 | 16047±1634 | 0.1616 | 0.1707 | 0.3 |
| | 8-HETE | 1564±141 | 1193±144 | 1307±163 | 1249±240 | 0.2571 | 0.5894 | 0.4035 |
| | 12-HETE | 9381±1843 | 9891±1508 | 11233±1514 | 8483±1497 | 0.5007 | 0.8929 | 0.3311 |
| | 15-HETE | 14501±1298 | 10808±2031 | 11537±989 | 10395±1148 | 0.0941 | 0.231 | 0.3605 |
| | 5oxoETE | 17366±1318 | 20500±3578 | 16570±1933 | 21987±8886 | 0.4158 | 0.947 | 0.8261 |
| | LxA4 | 1538±197 | 838±186 | 928±104 | 779±196 | *0.0287 | 0.0757 | 0.1367 |
| | LXB4 | 3134±478 | 3422±639 | 3308±355 | 3556±1006 | 0.697 | 0.8228 | 0.9774 |
| | 8isoPGA2 | 3238±549 | 2420±639 | 2441±424 | 2554±332 | 0.4794 | 0.5059 | 0.3541 |
| | PGD2 | 2738±439 | 1877±342 | 1993±443 | 1765±289 | 0.1851 | 0.292 | 0.4317 |
| | PGE2 | 2499±513 | 1835±409 | 1729±330 | 2104±137 | 0.7025 | 0.5121 | 0.1841 |
| | PGF2a | 24286±5329 | 24787±4117 | 28916±5965 | 24911±4527 | 0.7398 | 0.6528 | 0.6697 |

| | | | | | | | | |
|---|---|---|---|---|---|---|---|---|
| | 15dPGJ2 | 235±55 | 199±56 | 194±49 | 198±38 | 0.7557 | 0.6748 | 0.7021 |
| | 6kPGF1a | 2276±430 | 2074±495 | 3275±422 | 2165±220 | 0.1198 | 0.1911 | 0.2714 |
| | TXB2 | 5595±917 | 6473±984 | 4331±451 | 6341±2026 | 0.2749 | 0.5919 | 0.6631 |
| DHA | 14-HDoHE | 1797±282 | 346±47 | 1690±261 | 334±53 | ****<0.0001 | 0.7776 | 0.8216 |
| | 17-HDoHE | 1629±199 | 357±64 | 1530±202 | 380±80 | ****<0.0001 | 0.8165 | 0.7049 |

**Supplementary Table 10:** Hippocampal oxylipin profile at P21: Data are represented as mean ± SEM (pg/mg of protein). N = 5 mice/group. Statistical comparisons were performed for each oxylipin using a 2 way ANOVA (sex x diet). LA: linoleic acid; AA: arachidonic acid; DHA: docosahexaenoic acid.
